# The shared selection landscape of dog and human cancers

**DOI:** 10.1101/2025.10.15.682624

**Authors:** Diane P. Genereux, Kate Megquier, Ross Swofford, Jason Turner-Maier, Michelle White, Vista Sohrab, Christopher Husted, Heather Gardner, Corrie Painter, Cheryl London, Elinor K. Karlsson

**Affiliations:** Medical and Population Genetics Program, Broad Institute of MIT and Harvard, Cambridge, MA 02142, USA; Genomics and Computational Biology, University of Massachusetts Chan Medical School, Worcester, MA 01655, USA; Morningside Graduate School of Biomedical Sciences, University of Massachusetts Chan Medical School, Worcester, MA 01655, USA; Cummings School of Veterinary Medicine at Tufts University, North Grafton, MA; Precede Biosciences, Boston, MA; Darwin’s Ark Foundation, Seattle, WA 98026, USA; Program in Molecular Medicine, University of Massachusetts Chan Medical School, Worcester, MA 01655, USA

## Abstract

Cancers in pet dogs are prevalent, progress rapidly, and closely resemble human cancers, positioning them as powerful models for precision oncology. While genetic drivers of human cancer often transcend histologic boundaries, most comparative studies have focused on matched cancer types, leaving the broader scope of genomic similarity unresolved. We performed the first exome-wide, histology-agnostic comparison of canine and human cancers, analyzing 429 dog and 14,966 human tumors across 39 types. Mutational signatures and genes under selection are widely shared between species, and cancer types are as genomically similar between species as within species, with no greater similarity within dog breeds than between breeds. Machine-learning models identify genetic features shared by dog and human tumors of different histologies, mirroring cross-histology patterns in human cancer. These findings establish dog cancer as a powerful system for genomics-informed precision oncology and support pan-cancer approaches to discover translationally relevant models beyond histologic classification.

## Introduction

Genomics-informed precision medicine has yielded promising treatments by targeting somatic mutations in tumor cells^1–6^. Some therapies initially approved for one cancer type have later proven effective against other cancers with mutations in the same genes^7–9^. For example, vemurafenib was initially developed to treat melanomas with V600E mutations in the *BRAF* gene. Subsequent pan-cancer studies revealed that it is also effective against tumors of 13 other cancer types that have the *BRAF* mutation, including several for which targeted treatments were not yet available^7^. Identifying genomic alterations shared across cancer types can reveal new treatment options for rare cancers, for which therapeutic development is often limited by small patient populations^10^.

Even so, effective genomics-informed treatments remain unavailable for many patients. Existing drugs target mutations in only a small set of genes, and acquired resistance limits long-term efficacy^11,12^. Identifying targetable driver mutations amid the background of passenger mutations often requires large patient cohorts, which are difficult to assemble^13,14^. Once a promising target is found, drug development is slow and costly, and efficacy of small molecule inhibitors is not guaranteed. Clinical trials can span a decade and cost as much as $2.8 billion^15^, and only about 10% of candidate drugs are ultimately approved^16^. Barriers are even steeper for the so-called “long tail” of rare driver mutations that occur in <5% of tumors, making it difficult or impossible to assemble patient cohorts adequate for target discovery and clinical testing^17–19^.

Pet dogs offer a powerful and underutilized opportunity to accelerate development of targeted therapies in a widely accepted preclinical model with a very large patient population ^20–24^. An estimated four million dogs are diagnosed with cancer each year in the United States^25^. Many pet dogs receive sophisticated care at veterinary specialty hospitals that parallels best practices in human oncology, including imaging, surgery, chemotherapy, immunotherapy, and radiation^26^. Furthermore, the less restrictive regulatory frameworks in veterinary medicine mean that new therapies can be tested in treatment-naive tumors, and dogs’ faster disease progression expedites evaluation of outcomes. The unique population structure of pet dogs includes hundreds of genetically isolated subpopulations (breeds)^27,28^, some of which have markedly high cancer risk^29–32^.

Cancer in pet dogs recapitulates many key features of human disease that can affect drug response and disease outcome^33–35^. Like other mammals, dogs acquire somatic mutations at a rate inversely correlated with their lifespan, and an adult dog has an equivalent number of somatic mutations in normal tissues as an adult human^36^. Dogs encounter many of the same environmental and lifestyle risk factors as humans and develop genetically heterogeneous, spontaneous tumors within the context of an intact immune system^37–40^. Cancer in laboratory mice, in contrast, is typically induced through genetic manipulation in animals with suppressed or absent immune function, and even immunocompetent transgenic mouse models do not recapitulate the heterogeneity of human tumors^41,42^. Alternatives such as human cancer xenografts or humanized mice address some of these limitations, but still fail to capture the full complexity of an immunocompetent host^41,43^.

Despite its potential to accelerate the development of genomically-informed targeted therapeutics, the dog model remains underexplored. Most studies to date have chosen dog cancers as models for human cancers based on matching by tissue of origin or clinical features^44–54^. However, it is increasingly clear that dog and human tumors from different tissues can share genomic features, including some relevant to treatment response. Clinical trials of tyrosine kinase inhibitors in dogs with mast cell tumors informed therapies for human gastrointestinal stromal tumors, as both cancer types have *KIT* gene mutations^55–58^, and dog osteosarcomas often have mutations in the genes *SETD2* and *DMD*^59^, which are rarely mutated in human osteosarcoma but commonly mutated in other human cancers^60,61^. Leveraging shared genomic features between dogs and humans, including across different histologies, may help to inform how interactions among somatic mutations influence responses to targeted therapy. Dogs with *BRAF*-mutant urothelial carcinoma exhibit lower rates of clinical responses to the targeted agent vermurafenib^62^, and could potentially help to elucidate the mechanism behind treatment resistance in *BRAF*-mutant human cancers with lower response rates, such as colorectal carcinomas^63^.

Here, we comprehensively evaluate the genomic similarity of dog to human cancers, and assess the potential of dog cancers to reveal new models for developing genomics-informed targeted therapeutics.

## Results

### Constructing a comparative cancer dataset

We constructed the largest cross-species comparative cancer dataset to date by compiling publicly available somatic mutation data for human and dog tumor exomes. Each tumor sample included in our set had a paired sample from normal tissue, enabling accurate detection of somatic (non-germline) mutations. We considered exclusively exonic variants (both single nucleotide variants (SNVs) and short insertions/deletions) because these were the mutation calls available for most tumors (77% in humans and 82% in dogs)(**Table S1, Fig. S1**). In total, we included 14,966 human tumors of 32 different cancer types from the Cancer Genome Atlas (TCGA)^64^, International Cancer Genome Consortium (ICGC)^65^ and CBioPortal^66^, as well as 429 dog tumors of seven different types of cancer^59,67–73^ (**Table S2, Fig. 1A**). The number of human tumors of each cancer type ranges from 45 angiosarcomas to 1784 breast cancers (mean=468±413 tumors per cancer type). The number of dog tumors per cancer type ranges from 23 mammary tumors to 109 B-cell lymphomas (mean=61±31)

**Figure 1.**
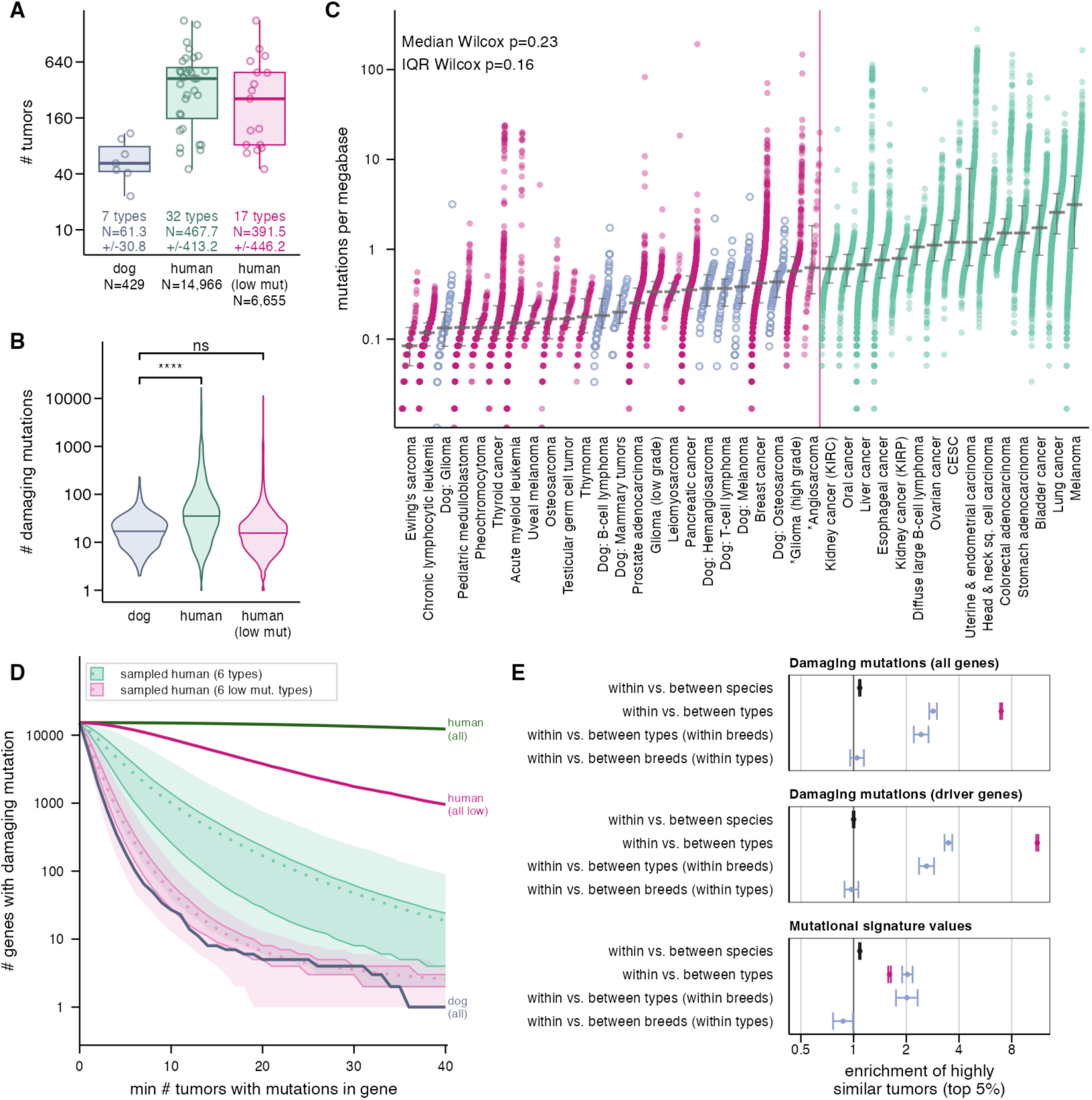
Differences between tumors are attributable to cancer type rather than species differences. **A.** Dataset includes 7 dog (blue) and 32 human (green) cancer types, including 17 we categorized as “low mutation rate” in our analysis (see panel C). Boxplots show median (line), IQR (box), and 1.5× IQR (whiskers); points represent individual tumors (jittered). **B.** Dog tumors harbor fewer damaging mutations than human tumors overall (p = 2.8×10⁻⁴⁷), but are not significantly different from low mutation rate human tumors (p = 0.19, two-sided Wilcoxon rank-sum). “Low mut” set excludes human cancer types with unusually high mutation rates. **C.** Mutation frequency per tumor, ordered by cancer-type median, modeled on prior work in human cancer^79^. Thick gray bars indicate medians; thin bars indicate IQRs. Species differences are not significant after accounting for cancer type. **D.** The higher number of damaging mutations in human cancers is explained by larger sample sizes and the high mutation rate cancer types. Downsampling to match dog sample sizes (1,000 permutations) shows comparable mutation frequencies between species. Dashed lines represent the median; the outer shaded ribbon the 95% range and the inner ribbon the 50% range. **E.** Tumors are no more similar within species than between species when considering either damaging mutations or mutational signatures, but are similar within cancer types. In dogs, tumors are not more similar within breeds. Colors: blue = dog; green = all human cancers; pink = low mutation rate human cancers.

For six of the seven dog cancer types, our dataset includes a human cancer type considered to be a histological match: osteosarcoma, melanoma, mammary tumors (breast cancer in humans), glioma (high-grade glioma in humans), B-cell lymphoma (diffuse large B-cell lymphoma in humans) and hemangiosarcoma (angiosarcoma in humans).

We analyzed 15,315 genes for damaging protein-coding mutations in dogs and humans. This included 15,289 genes with a single ortholog in dogs and 26 genes with two orthologs in dogs that are known cancer driver genes in humans (**Table S3**). In total, our list includes 1,098 genes known to drive cancer in humans (hereafter referred to as “driver genes”)^74–76^ and 4852 genes identified as potentially druggable targets by OncoKB^74^, including 464 driver genes (hereafter referred to as “druggable genes”). Most driver genes have not yet been functionally validated to also drive cancer in dogs. We classified as damaging all nonsynonymous mutations predicted to be damaging using either Ensembl Variant Effect Predictor or SnpEff^77,78^.

### Dog cancers model the genomic heterogeneity of human cancers

Overall, we found that cancer in dogs is a remarkably close genomic model for human cancers, with mutations in many of the same genes, often at the same genomic position, and a similar degree of genomic heterogeneity. We compared dog and human cancers in four dimensions: (i) the total number of mutations present; (ii) the distribution of mutations across the genome; (iii) the specific sets of genes mutated; and (iv) mutational signatures detected.

### Similar number of damaging mutations

The number of damaging mutations in dog versus human tumors is indistinguishable after accounting for cancer type. Overall, dog tumors have significantly fewer damaging mutations than human tumors (Fig. 1B). However, in both species, cancer type has a significant effect on mutation rate (dog ANOVA effect=0.11, F=8.7, p=6.9×10^−9^; human ANOVA effect=0.09, F=47.0, p=1.8×10^−273^). p=6.9×10^−9^; human ANOVA effect=0.09, F=47.0, p=1.8×10^−273^). When analyzed by cancer type, dog and human tumors have similar median mutation rates (p_wilcox_= 0.27) and have similar variability as measured by interquartile range (p_wilcox_=0.19) (Fig. 1C). This suggests that the higher overall mutation burden observed in the human dataset is explained by a greater representation of tumor types with higher mutation rates.

We defined a set of human cancer types with mutation rates comparable to those observed in dogs (“low mutation rate human cancer types”) by excluding nearly all human cancer types with a median mutation rates higher than the most highly mutated dog cancer type (dog osteosarcoma; median = 0.43 mutations/Mb) (Fig. 1C), with two exceptions. Both high-grade glioma (0.57 mutations/Mb) and angiosarcoma (0.62 mutations/Mb) had mutation rates slightly above our threshold (0.57 and 0.62 mutations/Mb respectively), but were also histiological counterparts for dog cancer types. We did exclude two histological matches that were among the most mutated human cancer types: diffuse large B-cell lymphoma (1.06 mutations/Mb) and melanoma (3.14 mutations/Mb). By retaining high-grade glioma and angiosarcoma, we were able to more robustly compare histologically matched and unmatched cancer types across species. Mutation rates in our “low mutation rate human cancer types” dataset are indistinguishable from dog cancers (Fig. 1B).

The rates at which genes are mutated is remarkably similar in dog and human tumors. Nearly all genes (15,313/15,315) were mutated in at least one of the 15,008 human tumors; a much smaller percentage of genes are mutated in at least one dog tumor (5462 genes; 36%; ks.test D=0.72, p=0). This difference was driven by the larger sample numbers and high mutation rate cancer types in humans, as it vanishes when we randomly downsample the human dataset to match the numbers of samples and cancer types in dogs, and exclude high mutation rate human cancer types (**Fig. 1D**)^79^. Two genes were not mutated in any human tumor: RPP21, which encodes a subunit of nuclear ribonuclease P and was also not mutated in dogs, and CD24, which encodes a cell adhesion protein and was mutated in a single dog tumor (a highly mutated melanoma).

### Genes mutated in dog and human tumors are largely the same

With regard to the specific genes mutated, dog and human tumors of different cancer types are as similar as human tumors of different cancer types. We quantified mutational similarity for pairings of tumors using cosine similarity, which ranges from 0 (none of the same genes mutated) to 1 (all the same genes mutated)(**Table S4**). After excluding highly mutated cancer types, we randomly sampled up to 200 tumors per cancer type, yielding 3,031,953 human/human pairs, 91,806 dog/dog pairs and 1,056,627 dog/human pairs. The similarity of dog/human pairs (mean=0.005) is comparable to the similarity of tumors from different cancer types within species (dog/dog mean=0.0061; human/human mean=0.0039)(**Table S5**; **Fig. S2**). Highly similar tumor pairs are only slightly more likely to come from the same species (odds ratio = 1.09; CI 1.07-1.10; p_fisher_=5×10-54), and this enrichment disappears when considering only driver genes (odds ratio = 1.0; CI 0.99-1.01; p=0.94) (**Fig. 1E**; **Table S6**).

The heterogeneity of mutated genes within and between cancer types is also similar in dogs compared to humans^80^. Tumors are more similar within a given cancer type than between cancer types in both dogs (mean=0.0172 vs. 0.0061; p_wilcox_=0) and in humans (mean = 0.0228 vs 0.0039; p_wilcox_=0). There is also a significant enrichment for highly similar tumor pairs within cancer types in both dogs (OR=2.8; CI 2.7-3.0; p_fisher_=0) and humans (OR=7.0; CI 6.9-7.1; p_fisher_=0)(**Fig. 1E**). The highest mean similarity for human tumors was between low-grade gliomas (median=0.056 for 19,900 pairs) and for dogs between pairs of hemangiosarcomas (mean=0.025 for 946 pairs)(**Table S7**, **Fig. S2**).

### Minimal effect of dog breed on spectrum of damaging mutations

We found no evidence that somatic mutations in tumors differ substantially between dog breeds. Our dataset includes dogs from over 50 dog breeds. When considering both cancer type and breed, breed had no significant effect on the number of mutations (Anova ges=0.047, p=0.31 for 16 breeds with three or more dogs; ges=0.051, p=0.004 for cancer type) (Fig. S3). There is no difference in the cosine similarity of tumors from different breeds compared to those from the same breed (p=0.06 for all genes and p=0.9 for driver genes)(Fig. S2). Tumors of the same type from the same breed are also no more enriched for highly similar pairs than tumors of the same type from different breeds when considering all genes (OR=1.05; p=0.33) or just driver genes (OR=0.97; p=0.55) (Fig. 1E).

### Genes shared between dogs and humans are enriched for driver genes

Genes with higher rates of mutation in both dog and human tumors are more enriched for driver genes than in either species independently, illustrating the potential for information from dogs to reveal genes relevant to cancer progression. We considered a gene mutated if it had one or more predicted-damaging mutations.

Overall, genes mutated in both human and dog tumors are enriched for both druggable genes (binomial p= 2×10^−4^, enrichment=1.1) and driver genes (binomial p = 1.9×10^−25^, enrichment=1.4). In both dogs and humans, enrichment for driver genes is most pronounced among the genes most frequently mutated (**Fig. 2A**). Even after correcting for gene-length, the fraction of tumors with mutations in a given gene is also correlated between the two species for genes overall (R^2^= 0.53; p=0), and especially for driver genes (R^2^=0.78; p= 4.2×10^−222^).

**Figure 2.**
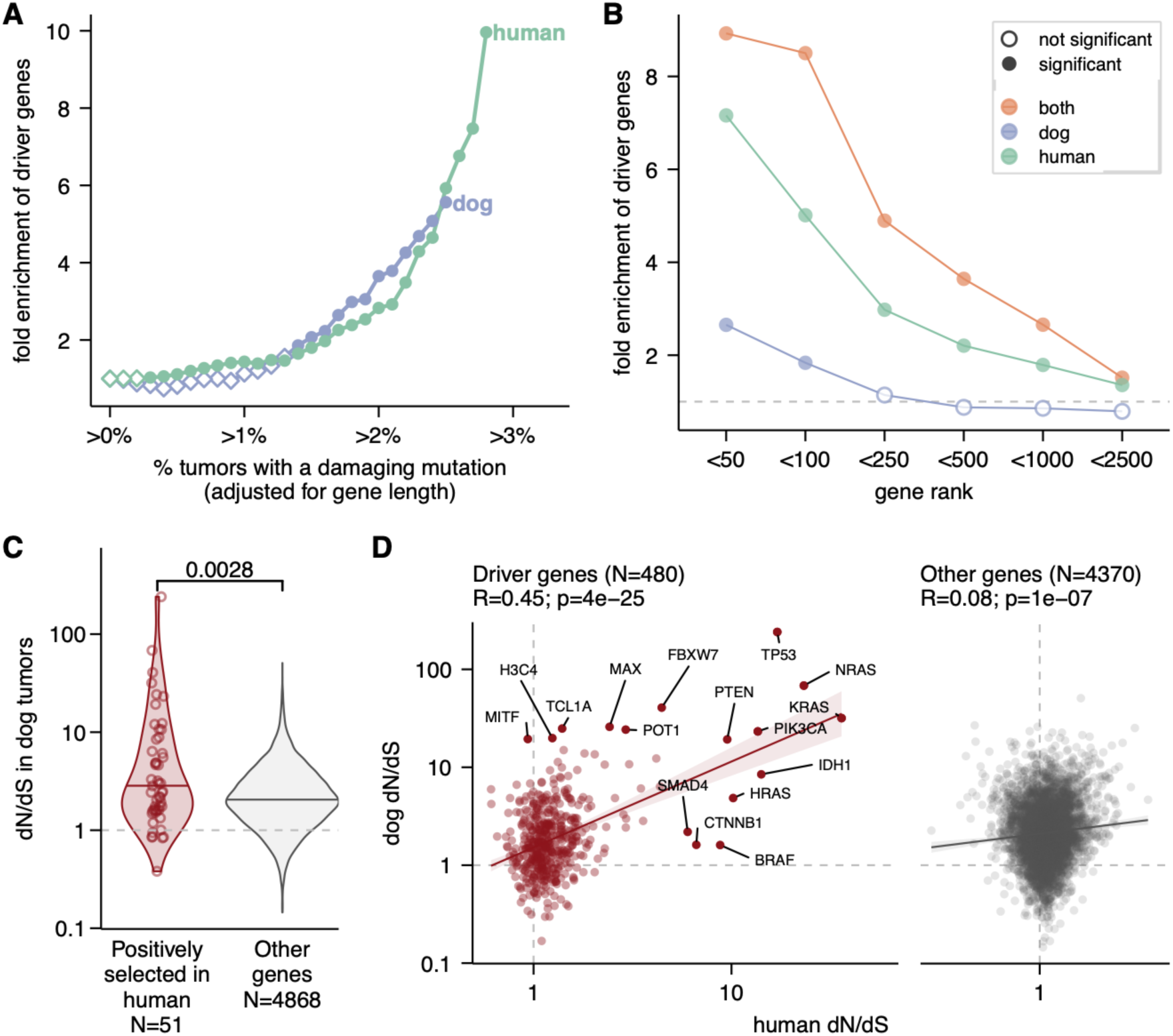
Driver genes are enriched for selection in both species. **A.** The most frequently mutated genes are strongly enriched for known driver genes, with a similar rate of increase in enrichment as the percentage of mutated tumors rises in both dog (blue) and human (green) tumors. **B.** Considering whether a gene has a high dN/dS in either species yields stronger enrichment for driver genes than analyzing each species independently. The dashed line indicates no enrichment. **C.** Genes under positive selection in human tumors (red; defined as having a dN/dS ratio for missense mutations > 1 and FDR-adjusted p-value < 0.05) show higher dN/dS values in dog tumors than other genes (gray). **D.** The dN/dS ratio for missense mutations is significantly correlated between dog and human tumors for driver genes (red), but not for other genes (gray). Dashed lines indicate dN/dS = 1. Linear regression lines are shown for each category, with shaded bands representing 95% confidence intervals. Pearson correlation coefficients (R) and corresponding p-values from two-sided correlation tests are shown above each plot. Genes ranked in the top ten in either species are labeled.

Our finding that the genes driving human cancer are particularly overrepresented among genes under positive selection in both dog and human tumors suggests that mutations in these genes confer a proliferative advantage in both species. Tests for positive selection can distinguish genes with mutations that confer growth advantage from passenger mutations^81,82^. For each gene, in each species, we calculated the ratio of nonsynonymous missense mutations to synonymous mutations (dN/dS) using the software dNdScv, which controls for variation in gene length and mutation rates^79^. Considering whether a gene has a high dN/dS in either species yields a stronger enrichment for driver genes than considering each species on its own (**Fig. 2B**). Across all genes, those with higher dN/dS score in human tumors tend to have higher dN/dS scores in dogs as well (**Fig. 2C**), and the strength of selection on each gene is correlated between the two species, especially for driver genes (**Fig. 2D**).

### Some driver genes rarely mutated in human tumors are commonly mutated in dogs

Some driver genes that are rarely mutated in human tumors are more commonly mutated in dog tumors, indicating potential for assembling larger study cohorts to evaluate targeted therapeutics. *POT1*, which drives angiosarcoma, a cancer rare in humans^83^, is mutated in 4.4% of the dog tumors in our set, compared to just 1% of all human tumors (p=4.8×10^−7^), *FBXW7*, a driver gene in uterine carcinosarcoma^84^, is mutated in approximately 6% of the dog tumors in our set, but only 1.9% of human tumors (p=5.8×10^−7^)(**Table S8**), and *DDX3X*, which is often mutated in B- and T-cell lymphoma and melanoma^85^, is mutated in ~4% of all dog tumors, but just ~1% of human tumors (p=7.7×10^−5^). The prevalence of these mutations in part reflects the large proportion of dog B-cell lymphomas (25.4% of all dog tumors) in our dataset. All three of these genes are under positive selection (dN/dS>1 and p_FDR_<=0.05) in dog B-cell lymphomas and frequently mutated (17.4%, 21.1%, and 12.8% for *POT1*, *FBXW7* and *DDX3X* respectively). In humans, mutation rates peak for these three genes at 17.8% for *POT1* in angiosarcomas, 17.8% for *FBXW7* in uterine & endometrial carcinomas and 6.2% for *DDX3X* in pediatric medulloblastomas (**Table S9**).

Many genes are under stronger selection in a dog cancer type than in any human cancer type. About 16.1% of all genes mutated in dog tumors (882 genes), and 11.9% of driver genes (67 genes), have higher dN/dS in at least one dog cancer type than in any of the 32 human cancer types we analyzed (**Table S9**). Fourteen genes are under significant selection (p_FDR_<0.05) in at least one dog cancer type, despite the relatively small sample numbers for each dog cancer type. These include *SATB1,* which is under selection in both human thyroid cancer (THCA) and dog T-cell lymphoma, but is rarely mutated in humans (frequency=0.006) and commonly mutated in dogs (freq=0.29). It also includes *PIK3CA,* which is mutated in 31.8% of dog hemangiosarcomas, a higher frequency than in any human cancer type except uterine & endometrial carcinomas.

### Mutation hotspots are shared

In addition to sharing many of the same frequently mutated genes, dog cancers often have mutations at the same frequently mutated “hotspot” positions within those genes (**Fig. 3A**; **Table S10**). We here define a site as a hotspot if it was either previously reported as a human hotspot^86,87^ and or mutated in five or more samples in our set.

**Figure 3.**
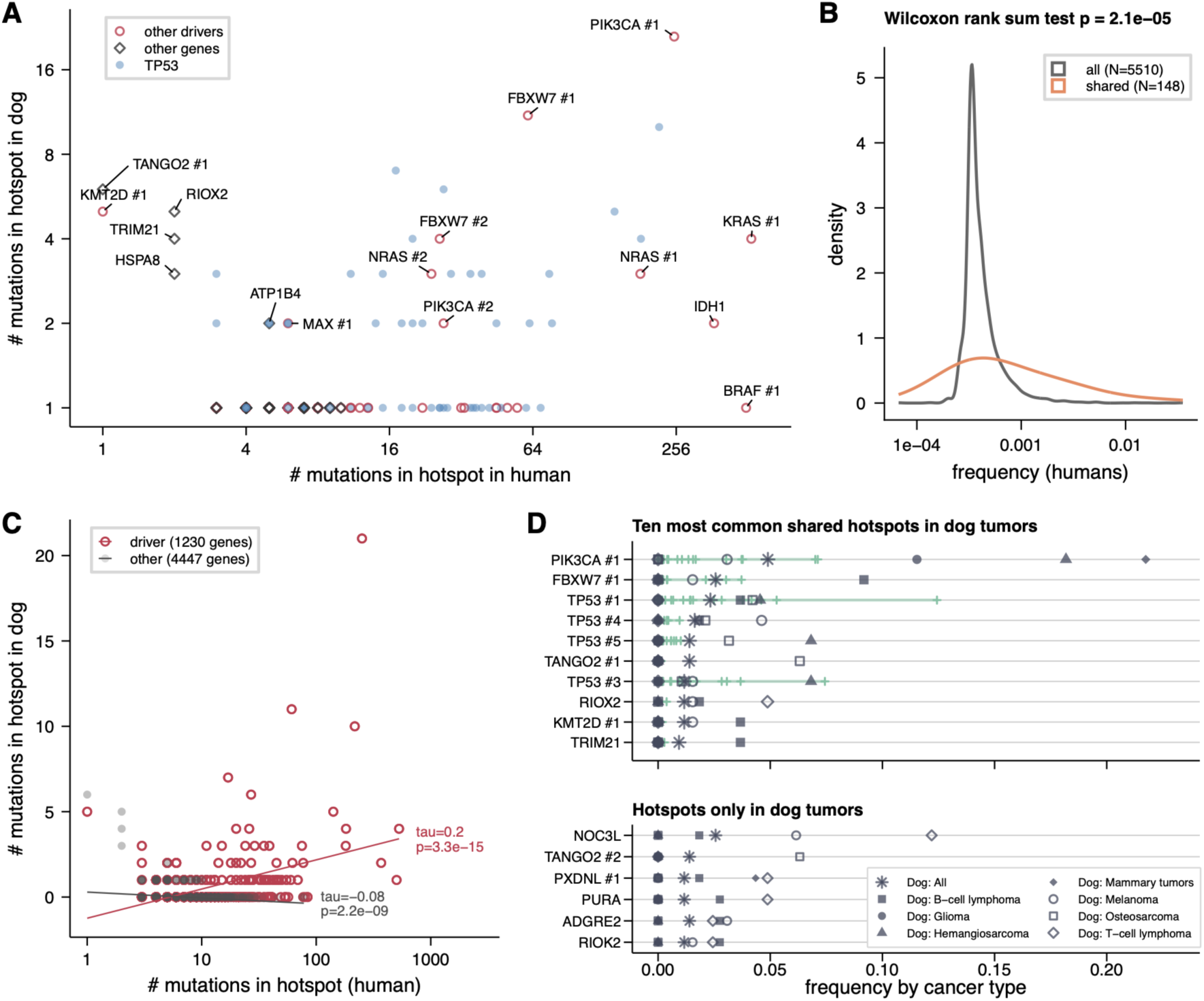
The same mutation hotspots are found in dog and human tumors. **A.** Nearly all mutation hotspots that are common in human tumors are also observed in at least one dog tumor, despite the much smaller number of dog tumors analyzed. Some mutations that are rare in humans appear more frequently in dogs. **B.** Shared hotspots tend to occur at significantly higher frequency in human tumors, suggesting that the smaller size of the dog dataset may limit detection of less frequent hotspots. **C.** For hotspots in driver genes (red), the number of mutations is significantly correlated between dogs and humans, while no such correlation is observed for hotspots in other genes (grey). The strength and significance of the association are reported as Kendall’s τ and the corresponding p-value. **D.** Of the ten most common hotspots in dogs that are also seen in human tumors, five occur at frequencies above 1% in at least one human cancer type. Six hotspots were observed in at least five dog tumors but in no human tumors. Dog cancer types are shown as grey symbols. The green horizontal line shows the range of frequencies across human cancer types, with each type indicated by a short vertical tick mark.

The five genes with the most commonly shared hotspots are driver genes: *TP53* (52/130 sites), *PIK3CA* (6/24 sites), *FBXW7* (6/12 sites), *PTEN* (3/24 sites) and *NRAS* (3/3 sites). This confirms previous work that described this phenomenon in smaller numbers of tumors^69,73^ and candidate genes^88^. We found 148 hotspots in humans that are also mutated in at least one of the 429 dog tumors. This is a relatively small percentage of all human hotspots identified from previous studies^87,89,90^ (2.7%; 148 out of 5377 hotspots in 3450 genes), but this is expected given the smaller size of the dog dataset. Nearly all human hotspots (99.1%; 5428 hotspots) have a frequency below our detection threshold in dogs (1/429 dogs). This suggests that if we had data for more dog tumors, and thus more power to detect lower frequency hotspots, we would discover more shared hotspots. Consistent with this, hotspots with higher frequencies in human tumors are more likely to also be found in dogs (p_wilcox_=1.9×10^−5^)(**Fig. 3B**).

Hotspots shared between dogs and humans are strongly enriched for cancer driver genes (hypergeometric p=5.9×10^−37^), and overall the frequency of hotspot mutations is correlated in driver genes but not other genes (**Fig. 3C**). Of the 10 most common hotspots in dogs, including some more common in specific dog cancer types, only four have a frequency greater than 0.02 in any human cancer type, and all four are in driver genes (**Fig. 3D**). Despite the smaller size of the dog dataset, we identified six “dog-specific” hotspots mutated in at least five dog tumors and no human tumors. This includes a hotspot in *NOC3L*, a gene involved in the initiation of DNA replication, found in 11 dog tumors, including 5 out of 41 T-cell lymphomas. Another dog-specific hotspot alters the adhesion G-protein coupled receptor *ADGRE* and is found in tumors from three different dog cancer types (B-cell lymphoma, T-cell lymphoma and melanoma). In humans, aberrant expression of *ADGRE* is associated with poor outcomes in patients with chronic myeloid leukemia^91^.

### Mutational signatures are shared

Dog and human tumors exhibit the same mutational signatures and at similar frequencies, suggesting that the biological processes, environmental exposures, and DNA repair deficiencies underlying cancer development are also shared. We screened each sample for all of the 43 mutational signatures detected in at least one tumor in our dataset. Within both dog and human cancer types, we find heterogeneity in the abundance of each signature (**Fig. S4, Fig. S5**). We considered a signature to be present in a sample if its value exceeded 0.05, consistent with published guidelines^92^(**Table S11**). Twenty-eight of the 43 signatures detected in human tumors were also detected in at least one dog tumor. The fraction of tumors with a given mutational signature was correlated between dogs and humans (Spearman’s rho=0.61; p=1.2×10^−5^)(**Fig. 4A**).

**Figure 4.**
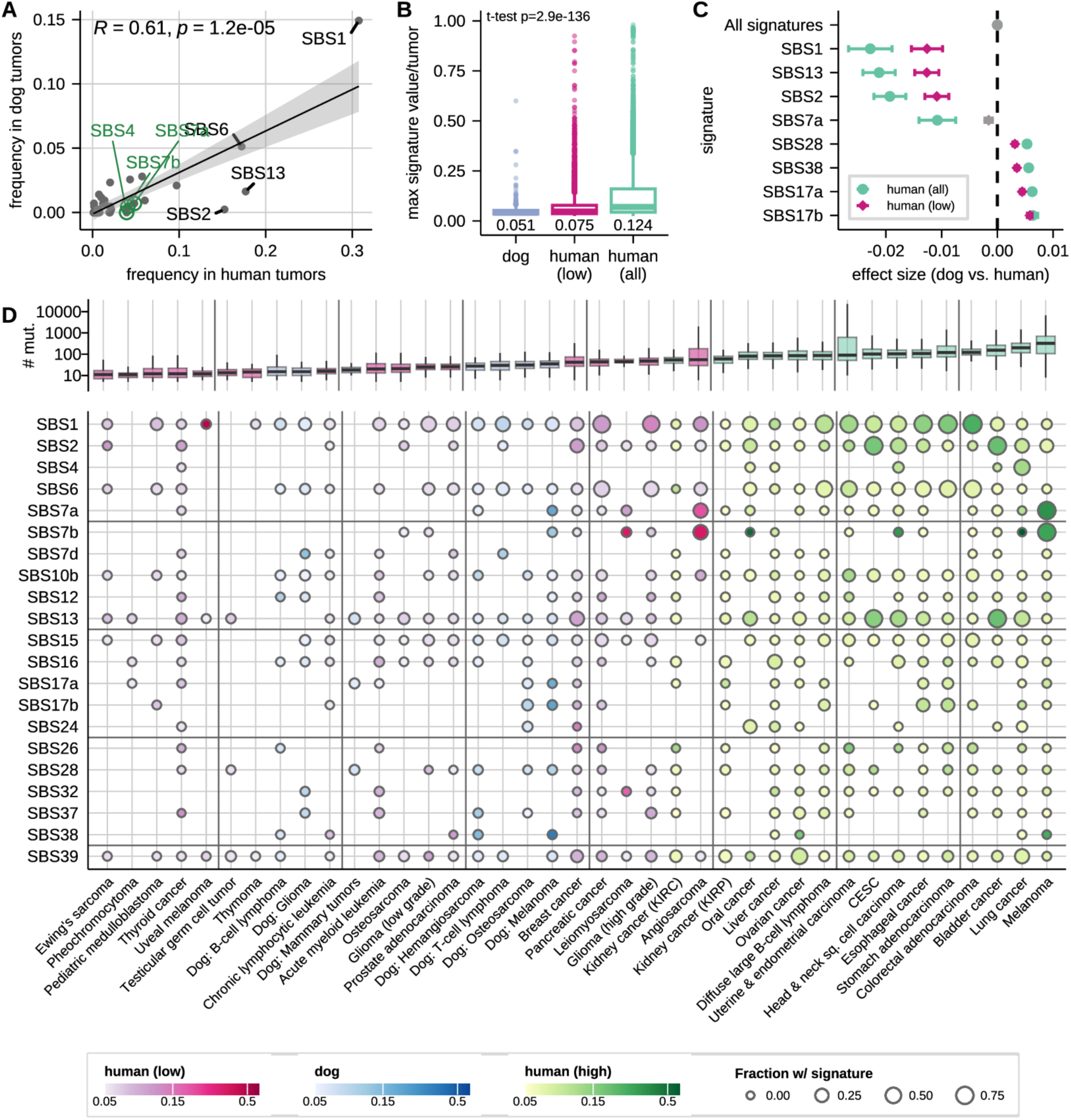
The same mutational signatures are found at similar frequencies in dog and human tumors. **A.** Correlation of mutational signature frequency between dog tumors and human tumors. The four most common signatures in human tumors are labeled in grey. Signatures linked to environmental exposures are labeled in green and occur at somewhat lower frequencies in dogs. These include SBS4 (tobacco smoke) and SBS7a and SBS7b (ultraviolet light). **B.** Human tumors show higher maximum signature values than dog tumors overall, but are more similar when only low mutation rate human cancer types are considered. Boxplots show the median (center line), interquartile range (box), and whiskers extending to 1.5× the interquartile range. Outliers are shown as individual points. **C.** Overall, species has no effect on signature values, but some signatures tend to be higher in humans (negative effect size) or higher in dogs (positive effect size). Non-significant (p_FDR_<0.05) effect sizes in gray. **D.** Mutational signatures detected in dog and human tumors. The most common signatures in dogs, and those in panel C, are shown here. Plot of all signatures shown in Fig. S6.

Seven of the top ten most common mutational signatures were shared between the two species. The aging-associated signature SBS1 was the most common in both, appearing in 31% of human and 15% of dog tumors. Three other signatures were present in at least ten dog tumors: SBS6 (17% in humans, 5% in dogs) and SBS15 (6% in humans, 3% in dogs), both implicated in defective DNA mismatch repair, and SBS10b, which is linked to defective proofreading activity of POLE (4% in humans, 3% in dogs)(**Fig. 4A**).

Overall, mutational signature values are broadly similar between dogs and humans. Although the maximum signature value per tumor is often higher in human tumors (**Fig. 4B**), linear regression indicates no overall effect of species (p = 1) or mutation count (p = 1) on signature values. However, there is a strong interaction between species and specific signatures (p < 2.5×10^−214^), explaining 9.1% of the total variance in signature values (**Fig. 4C**). The species effect varies by signature: values are significantly higher in dogs for 25 signatures (FDR < 0.05), including four with effect sizes >0.005, and significantly higher in humans for six signatures, also including four with effect sizes >0.005. For example, SBS1, associated with spontaneous deamination of 5-methylcytosine and cellular aging, is higher in humans, possibly reflecting their longer lifespan^93^. SBS2 and SBS13, linked to APOBEC activity, are also more abundant in human tumors^94^.

The mutational signatures detected in dog tumors suggests many, but not all, environmental risk factors for cancer are shared between dogs and humans (**Fig. 4D**). UV-associated signatures like SBS7a are common in humans but rare in dogs (detected in fewer than 1% of dog tumors), consistent with the protective effect of fur^95^. In turn, dog tumors tend to have higher values for SBS38, which is linked to indirect damage from UV-light^96^(**Fig. 4C**). However, when UV-associated signatures are detected in dogs, it is in cancer types linked to UV exposure in humans. The two human cancer types where SBS7a signatures are most common are melanoma (80%) and angiosarcoma (36%). In dogs, we detect SBS7a in cancers from two tissues not typically exposed to UV,oral melanoma and visceral hemangiosarcoma, the histological analog of angiosarcoma^97,98^, a phenomenon previously described in human lung melanomas^99^. We do not detect SBS4, a signature primarily associated with the human activity of tobacco smoking, in any dog tumors, although our dataset does not include dog nasal carcinomas, which have been associated with second-hand smoke exposure^100^. In four dog osteosarcomas, we detect SBS24, a signature proposed to be linked in humans to exposure to the mycotoxin aflatoxin^101^, a known possible contaminant in dry pet food^102^. We also find signatures associated with exposure to the carcinogenic herb *Aristolochia* in humans and mice^103,104^ (SBS22 in one dog B-cell lymphoma) and with prior treatment with azathioprine in humans (SBS32 in one dog glioma)^105^. Azathioprine is a purine anti-metabolite also used to treat autoimmune diseases in dogs^106^. We note, though, that while mutational signatures are known to be shared across mammals^36^, the associations of specific signatures with environmental and endogenous exposures have primarily been discovered in humans^96^ and have yet to be validated in dogs.

### Dog cancers resemble human cancers in variation of mutational signatures present

Dog tumors are almost indistinguishable from human tumors in terms of mutational signature values once the high mutation rate human cancer types are excluded. We calculated cosine similarity for all tumor pairs (3,031,953 human/human pairs, 91,806 dog/dog pairs and 1,056,627 dog/human pairs). The median similarity of tumors of different types was similar in dogs (median=0.955 +/− IQR 0.053; N_pairs_=76,030), humans (median=0.950 +/− IQR 0.117, N_pairs_=2,823,437) and when comparing dog and human tumors (median=0.953 +/− IQR 0.075; N_pairs_=1,056,627) (Table S5).

Tumors pairs from the same species are slightly enriched for close matches (OR=1.09; CI 1.07-1.10; p_fisher_=5×10^−55^), but much less so than same-species tumor pairs from the same cancer type in both dogs (OR=2.03; CI 1.89-2.17; p_fisher_=4×10^−84^) and humans (OR= 1.60; CI 1.58-1.63; p_fisher_=0) (**Fig. 1E**). In dogs, breed has little effect. Tumors of the same type from the same breed have a slightly lower proportion of high-similarity pairs than do tumors of the same type from different breeds (OR=0.870; CI 0.763 - 0.991; p_fisher_=0.03), suggesting that dog’s breed does not strongly influence the mutational signatures in the tumor (Table S7).

### Matching dog and human tumors with high genomic similarity

#### Unsupervised clustering finds no distinct clusters

No dog cancer type is distinct from human tumors or closely aligned to any single human cancer type when tumor relationships are visualized using t-distributed Stochastic Neighbor Embedding (t-SNE). To cluster based on mutational signatures, we ran t-SNE on the principal components that together explained more than 70% of the variance (**Table S12**, **Table S13**). To cluster based on driver gene mutations, we applied t-SNE directly to the matrix of mutated driver gene counts. We used this approach because the mutation matrix is sparse, and each principal component explains only a small fraction of the variance, even using methods optimized for sparse data. Consistent with our comparisons of driver mutations and signatures in dog and human cancers, dog tumors overlap low mutation rate human cancer types in the t-SNE analysis (**Fig. S7**). The lack of separation of dog tumors from low mutation rate human tumors is even more evident in a t-SNE analysis that excludes the high mutation rate human cancer types (**Fig. 5AB; Fig. S8; Fig. S9**).

**Figure 5.**
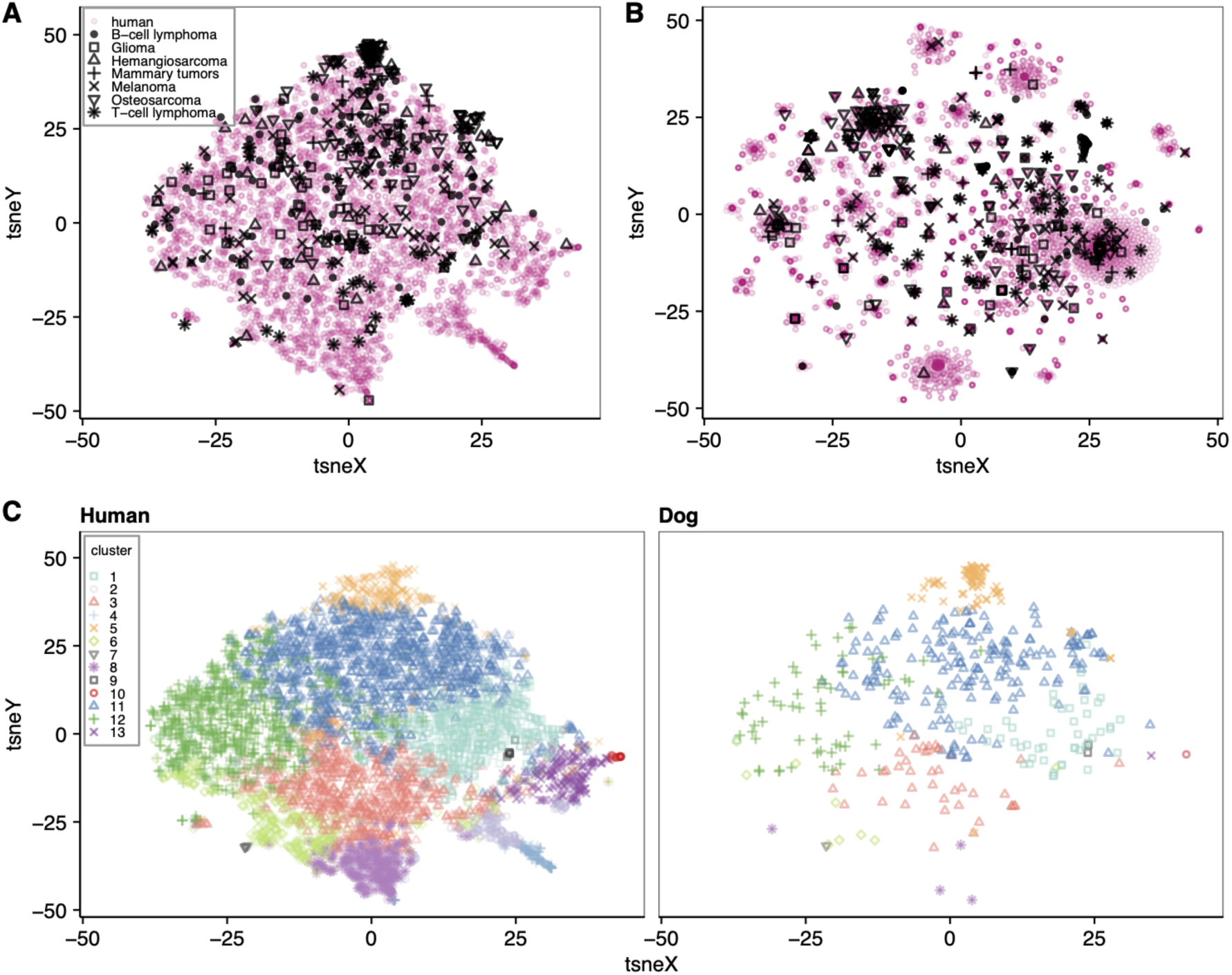
Unsupervised clustering does not separate dog cancers from human cancers. Dog cancers overlap low mutation rate human cancers and do not separate by cancer type in t-SNE visualizations based on **(A)** principle components explaining 70% of sample variance in mutational signatures and **(B)** which driver genes are mutated. (**C)** K-means clustering using the mutational signature principal components yields 13 clusters, 11 of which include both human and dog tumors.

Clustering tumors using the mutational signature principal components similarly showed that dog tumors are intermingled with human tumors. We performed k-means clustering and selected the number of clusters using the elbow method based on the within-cluster sum of squares. When clustering all tumors (10 clusters), 94% of dog tumors were assigned to the largest single cluster, together with half of all human tumors (**Fig. S10A,B**). This proportion decreased to 43% when we included only low mutation rate human tumors (13 clusters), and the number of clusters containing ten or more dog tumors increased from two to five (**Fig. 5C**; **Fig. S10B**). Each dog cancer type was distributed across at least four clusters, and clusters with a more heterogeneous representation of human cancer types, as measured by the Shannon entropy, were also more heterogeneous in dogs (**Fig. S10C**)(Spearman rho=0.93, p=4×10^−5^). We observed some enrichment of certain cancer types within specific clusters. For example, both dog osteosarcomas (observed = 50, expected = 16.6, p_FDR_ = 7×10^−19^) and human thymomas (observed = 13, expected = 3.6, p_FDR_ = 0.009) were enriched in cluster 5 (**Fig. S10D**).

#### Supervised machine learning finds cross-species genomic similarities between cancer types

We used supervised machine learning to identify pairs of dog and human cancer types enriched for shared genetic features, even in the absence of broader genomic similarity. Our approach, which was previously used to classify human cancers^107^, could identify dog cancer types that are potential models for trialling targeted therapies. We first learn the genomic features of the histology-labeled human tumors (our training set), then use those features to classify each dog tumor as a human cancer type. To quantify the relative contribution of individual features to classification, we computed Shapley values^108,109^. This approach is well-suited for classifying samples using genomic features common in the human cancer types well-represented in our data set. It may miss pairs previously reported to share genomic features if there are few human samples in the training set or if the shared genomic features are not included in our model (e.g. noncoding, structural, and epigenetic changes^70^). It will also miss histology-matches with few shared genomic features because of differing etiology (e.g. angiosarcoma and melanoma in humans frequently occur in sun exposed skin).

To implement machine learning, we use eXtreme Gradient Boosting (XGBoost^110^) models with up to 40,189 features, including mutated genes and pathways, mutational signature values, and overall mutation count. XGBoost is a decision tree–based algorithm that is well-suited for somatic mutation data, as it efficiently models non-linear relationships and interactions among sparse, high-dimensional features while filtering out uninformative variables. We trained models on two groups of human cancers: (1) 17 low mutation rate cancer types and (2) all cancer types. For each group, we evaluated three different feature sets: the “full model” (655 features retained as informative by machine learning), a “driver gene mutations model” (all 1098 driver genes), and a “mutational signatures model” (all 43 signatures). We calculated the average accuracy of the classifier on held-out human data across 5 folds using the k-fold cross validation procedure^111^. Using the full model, tumor classification accuracy was 63.5% for low mutation rate cancer types and 63.4% for all cancer types (**Fig. S11**, **Fig. S12, Table S14**). This is comparable to previous classifiers trained on somatic mutations in exome sequencing data^112,113^.

We also assessed the performance of a model trained on the much smaller dataset of dog cancers. Accuracy under our dog-trained model was slightly lower (57.7%) than under our human-trained model (**Table S14, Fig. S11C**, **Fig. S12**). Classification accuracy for individual dog cancer types was positively correlated with the number of tumors of that type in the dataset (R=0.87; p = 0.018). Variation in classification accuracy across cancer types was explained in part by the number of samples available (11% of variance explained; p=0.047), and not by species (0% of variance explained; p=0.99).

#### Classifying dog tumors as human cancer types

We applied the human-trained classifier to assign each dog tumor to a human cancer type, based on genomic features. Using models trained on low mutation rate cancer types, tumors of five of the seven dog cancer types were assigned to fewer human cancer types than expected by chance. We quantified this phenomenon of “clumpiness” using Shannon’s entropy, and measured the statistical significance via permutation (**Table S14**). Of the seven dog cancer types, hemangiosarcoma, osteosarcoma and T-cell lymphoma are clumpy in both the full model and at least one other model (**Fig. 6A**). Glioma and B-cell lymphoma are clumpy only in the mutational signatures model. Using models trained on all cancer types, six of the seven cancer types were clumpy in at least one model (**Fig. S13A**).

**Figure 6.**
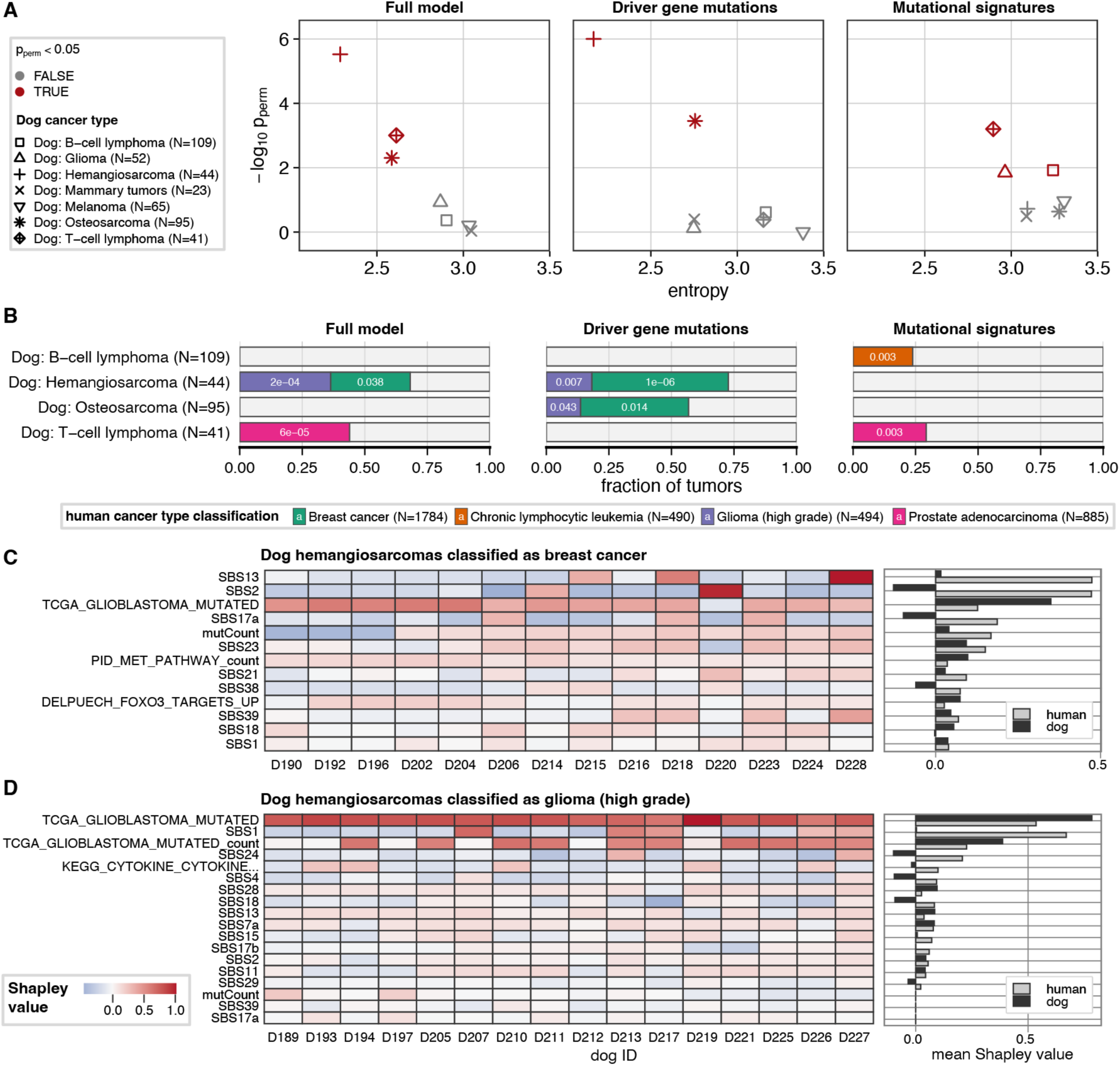
Supervised machine learning finds significant matches between human and dog cancer types using genomic features. **A.** Dog cancers in red were classified as fewer human cancer types than expected by chance, suggesting enrichment for shared genomic features. **B.** A subset of dog cancer types were classified to specific human cancer types more than expected by chance, and this varied depending on the type of genomic features included in the model. **C.** The genomic features explaining the classification of dog hemangiosarcomas as either breast cancer or **D.** high grade glioma varies between tumor samples. The label for the KEGG Cytokine–Cytokine Receptor Interaction pathway is truncated.

Dog tumors of all types were classified to multiple different human cancer types, consistent with the genomic heterogeneity of both human and dog cancer types. Even so, some dog cancer types were classified as a specific human cancer type more often than expected by chance (**Fig. 6B**, **Fig. S13B, Table S15)**. For example, using the full model trained on low mutation-rate cancer types, dog hemangiosarcoma samples were often classified as high-grade glioma (p=2×10^−4^) or breast cancer (p=0.038), and dog T-cell lymphoma was classified as prostate adenocarcinoma (p=6×10^−5^). Under all three models, tumor classifications were informed by multiple features.

Low sample counts likely limited classification of some dog tumors to histology-matched human cancer types. Dog tumors were more often assigned to human cancers that were better represented in the training set (Pearson’s r=0.49; p=1.6×10^−8^ for the full model). Histology-matched cancer types were among those with fewest samples, including B-cell lymphoma (N = 172), osteosarcoma (N = 117), and angiosarcoma (N = 45), which may explain why machine learning did not assign dog tumors to these cancer types any more than expected by chance, even when the dog cancer type was previously reported to share genomic features (e.g. diffuse large B-cell lymphoma^67^ (N=172), osteosarcoma^59,68^ (N=117), and angiosarcoma^45,69^(N=45)). In contrast, the histology-matched type for canine mammary tumors, human breast cancer, was the most abundant cancer type in the dataset (N = 1,784), and canine mammary tumors were assigned to this type at a frequency higher than expected by chance in the driver-only model (4 of 23, p = 0.0218), consistent with previous reports^73^.

The classification of most dog hemangiosarcomas as both breast cancer (14/44) and high-grade glioma (16/44) was informed by a combination of genomic features, illustrating the capacity of machine learning to integrate diverse signals. Classification of most hemangiosarcomas as either glioma or breast cancer was informed by mutations in the TCGA Glioblastoma Mutated^114^ gene set (**Fig. 6C-D, Fig. S14, Table S16**, **Table S17**), consistent with the established role of the PI3K pathway, which includes many of these genes, in these cancer types^45,69,115,116^. Classification as breast cancer versus high-grade glioma depended in part on which genes were mutated. *TP53* was mutated in every hemangiosarcoma classified as high-grade glioma (N=16/16, mean Shapley = 0.784), and multiple genes in this pathway were also mutated in seven tumors (mean subset Shapley=0.599). Mutations in the TCGA_Glioblastoma_Mutated gene set were present in most hemangiosarcomas classified as breast cancer (mean Shapley value in N=13/14 samples where this feature was positive (mean subset Shapley) = 0.384), most commonly *PIK3CA* (N=11/14). However, this classification was also informed by the APOBEC signatures in a subset of samples SBS13 (N=3/14, mean subset Shapley=0.548) and SBS2 (N=2/14, mean subset Shapley=0.557), with SBS13 being the most important feature in a single sample. APOBEC signatures are found in more than 75% of human breast cancers and promote metastasis and treatment resistance^117–119^.

In contrast, the classification of 18 out of 41 dog T-cell lymphoma as prostate adenocarcinoma was informed primarily by mutational signatures. For all 18 tumors, signatures were among the top explanatory features. Multiple signatures contributed to each classification. The only non-signature feature informative for most samples was low to moderate mutational burden (mean_dog_=34 mut/Mb) (N=17/18, mean subset Shapley =0.313), while the signatures informative for most samples included higher levels of SBS28, which is linked to decreased DNA proofreading by polymerase epsilon^120^ (N=17, mean subset Shapley = 0.386), the mismatch repair deficiency signatures SBS6 (N=16/18, mean subset Shapley =0.278) and SBS15 (N=16/18, mean subset Shapley=0.103), and the aging signature SBS1 (N=17/18, mean subset Shapley =0.198). In addition, a subset of classifications were informed by higher levels of UV light signature SBS7d^121^ (N=6, mean subset Shapley = 0.056) and lower levels of signatures related to reactive oxygen species^122,123^ (SBS17a, N=10, mean subset Shapley = 0.145) and alkylating agents^124^ (SBS11, N=18, mean subset Shapley = 0.057).

## DISCUSSION

Cancer is unequivocally the same phenomenon in dogs and humans. Genomic similarities exist across several dimensions: the number and frequency of mutations, the specific genes and hotspots mutated, and the mutational signatures present. Together these traits confer similarities so extensive that pairs of dog and human tumors are, on average, no more genomically distinct than pairs of samples of different cancer types within each species. The strength of positive selection on individual genes is correlated between the two species, suggesting that mutation sharing is not merely a byproduct of similar mutation rates for orthologous genes, but instead a reflection of cancer mechanisms shared between dogs and humans. Though some human cancer types, particularly those induced by high carcinogen exposure, have higher mean mutation counts and concomitantly higher mutational signature values than observed in any dog cancer, mutation counts for dog cancer types fall within the overall range for human cancers.

The similarity of somatic mutational landscapes in dog and human cancers is consistent with the deep conservation of canonical cancer pathways across eukaryotes. Dogs and humans are mammalian species that diverged approximately 94 million years ago^125^, with total evolutionary branch length of 1.99 substitutions per site^126^. The distance is only slightly greater than the distance of house mice (*Mus musculus*) to human (1.64 substitutions per site; about 87 million years). In contrast, most canonical cancer pathways, including those involved in DNA damage response and repair, are conserved across vertebrates (588 million years), and often across all eukaryotes (1598 million years)^127^. While some species may have evolved lineage-specific modifications to suppress cancer development^128^, a comparison of 16 mammalian species found that somatic mutation rates in normal intestinal tissue across mammals are evolutionarily constrained and inversely correlated with lifespan (thus, an old dog is comparable to an old human), and result from common mutational processes that produce shared mutational signatures^36^.

This positions dogs as a powerful natural model for development of new genomics-informed targeted cancer therapies. Unlike induced cancer models in mouse, for example, cancer in dogs is as genomically heterogeneous as in humans, with mutational landscapes that mirror human cancers. Some driver mutations that are rare in human cancers are common in dogs, offering a unique opportunity to develop therapeutics for genomically defined subsets of human patients^129^. Dog and human cancers also share mutational signatures, many of which have yet to be conclusively connected to a specific exposure or genetic risk feature. Dogs could therefore help connect environmental exposures to cancer risk, especially given their short lifespans and accelerated disease course.

Pet dogs could be particularly useful for investigating tumor evolution during patient treatment^130^. In dogs, new therapeutics can be more easily trialed in treatment-naive tumors, and longitudinal studies completed more quickly because of the faster disease course^131–133^. Our comparisons show that the strength of selection acting on cancer driver genes is correlated between dogs and humans, and that dog tumors may have mutational signatures associated with prior treatment or treatment resistance. Trials in dogs would support the development of approaches to improve accuracy in predicting therapeutic resistance, tumor progression, and recurrence^134^.

Our machine learning results demonstrate the value of dog cancers as models for precision oncology. For example, we find that dog hemangiosarcomas commonly share genomic features with two different cancer types in humans, breast cancer and high-grade glioma, including mutations in *PIK3CA* and *TP53*, respectively. Given that hemangiosarcoma is estimated to affect up to 300,000 pet dogs annually in the United States, with few effective treatment options available, studies in dogs with hemangiosarcoma could address the recognized need for a model to discern how *PIK3CA* mutations impact vascular cancer^135,136^. Similarly, shared mutational signatures between dog T-cell lymphoma and human prostate adenocarcinoma suggest shared mutagenic pressures. However, genomic heterogeneity within dog cancer types makes tumor histology alone an unreliable predictor of which dogs are most appropriate for inclusion in clinical trials of targeted therapies.

Our observation of genomic heterogeneity within histologically-defined cancer types of both dog and human aligns with the well-established phenomenon of molecular cancer subtypes within^137,138^ and across^139^ cancer types. This heterogeneity likely affected the performance of our machine-learning models, for which training data are labeled only by cancer type, and not by subtype. Mislabeled or ambiguous data can lead the model to learn incorrect patterns, thereby reducing predictive power and increasing error rates^140^. Unfortunately, while subtype annotations are available from some of the data sources used here, subtype labels are difficult to assign consistently across data sources^141^. Furthermore, for cancer types with relatively few samples, subtype labeling is often less reliable^137^. Subdivision reduces sample sizes even more, undermining model performance^142^. As datasets grow and standardized subtype annotations are widely adopted, incorporating them into cross-species machine learning approaches could reveal genomic similarities between dog and human subtypes missed when labelling only by cancer type and potentially identify genomic models for human cancer subtypes that currently lack animal models^135^.

Somewhat unexpectedly, we find that, while the prevalence of specific cancer types differs across dog breeds, the somatic mutational profile of each cancer type does not. This is in contrast to earlier studies in smaller cohorts^67,143^, and has important implications for the design of comparative oncology studies. For studies targeting a specific cancer type, focusing recruitment on breeds with higher prevalence will yield a study population with a more uniform germline background, potentially reducing confounding factors common in human trials, while still capturing relevant somatic mutation heterogeneity. For many studies, though, particularly those focused on specific somatic mutations, restricting by breed is unnecessary. This will enable the recruitment of larger, genetically diverse cohorts that better mirror the diversity of human patient populations.

Our study focused on single base substitutions and short insertions/deletions in coding regions, the variant types for which data were most commonly available for both dog and human. This may have contributed to why, for example, dog osteosarcomas were not more often classified as human osteosarcoma. In both species, osteosarcoma has an unusually high burden of somatic copy-number alterations and structural rearrangements^59,68,144–149^ that, in humans, have been used to define molecular subtypes with clinical relevance ^150,151^. Our approach also does not capture non-coding variants that can activate oncogenes, as with *TERT* promoter mutations in human melanoma^152,153^. Although driver mutations in non-coding genes and regulatory sequence are likely less frequent than in protein coding genes, more continue to be discovered as human whole-genome cancer datasets grow^154^. Regulatory elements are often conserved across placental mammals^155,156^, and mutations in conserved regulatory elements may affect activity of driver genes^157^. Extending machine learning training data to consider structural variants and non-coding mutations could identify additional dog cancer types as useful genomic models among, potentially expanding cohorts for pre-clinical testing of targeted therapeutics^158^.

Achieving this will require expanding the scale and scope of dog cancer genomic data. Human pan-cancer discoveries have been enabled by major collaborative efforts to share genomic data for thousands of patients. Here, we leveraged TCGA^64^, ICGC^65^ and CBioPortal^66^ to include data for 14,966 tumors from human patients. Despite dogs having a nearly two-fold higher annual cancer incidence than humans^159^, with approximately 4 million diagnosed cases per year in the United States^25^, available genomic data for dogs is very limited, even for just coding regions. Assembling exome data for the 429 dog tumors we consider here required aggregating results across eight different studies^59,67–73^.

Dogs represent a unique opportunity for translational cancer research, as veterinary oncology routinely uses many of the same drugs, treatment modalities, and clinical metrics as human oncology. Dog owners are highly engaged partners, motivated by past experiences, the hope for better care, and the opportunity to contribute to medical advances. Today, mortality rates for dogs remain high—even with treatment: hemangiosarcoma kills 90% of dogs within a year^160^; osteosarcoma has a <20% two-year survival rate^161^; oral malignant melanoma has a one-year survival of 30%^162^; and median survival times are 6.5 months for high-grade glioma^163^, 6–8 months for T-cell lymphoma, and ~12 months for B-cell lymphoma^164^. Establishing genomics as a routine and clinically valuable tool in veterinary oncology could rapidly expand available datasets, especially if data is shared through platforms like the NCI’s Integrated Canine Data Commons^165^, and provide patient populations for preclinical trials. By integrating dog cancer genomics into translational research pipelines, we can accelerate the development of targeted therapies for both human and veterinary oncology.

## Resource Availability

## Acknowledgements

The results published or shown here are in part based upon data generated by the TCGA Research Network: https://www.cancer.gov/tcga.

---

**Figure S1.**
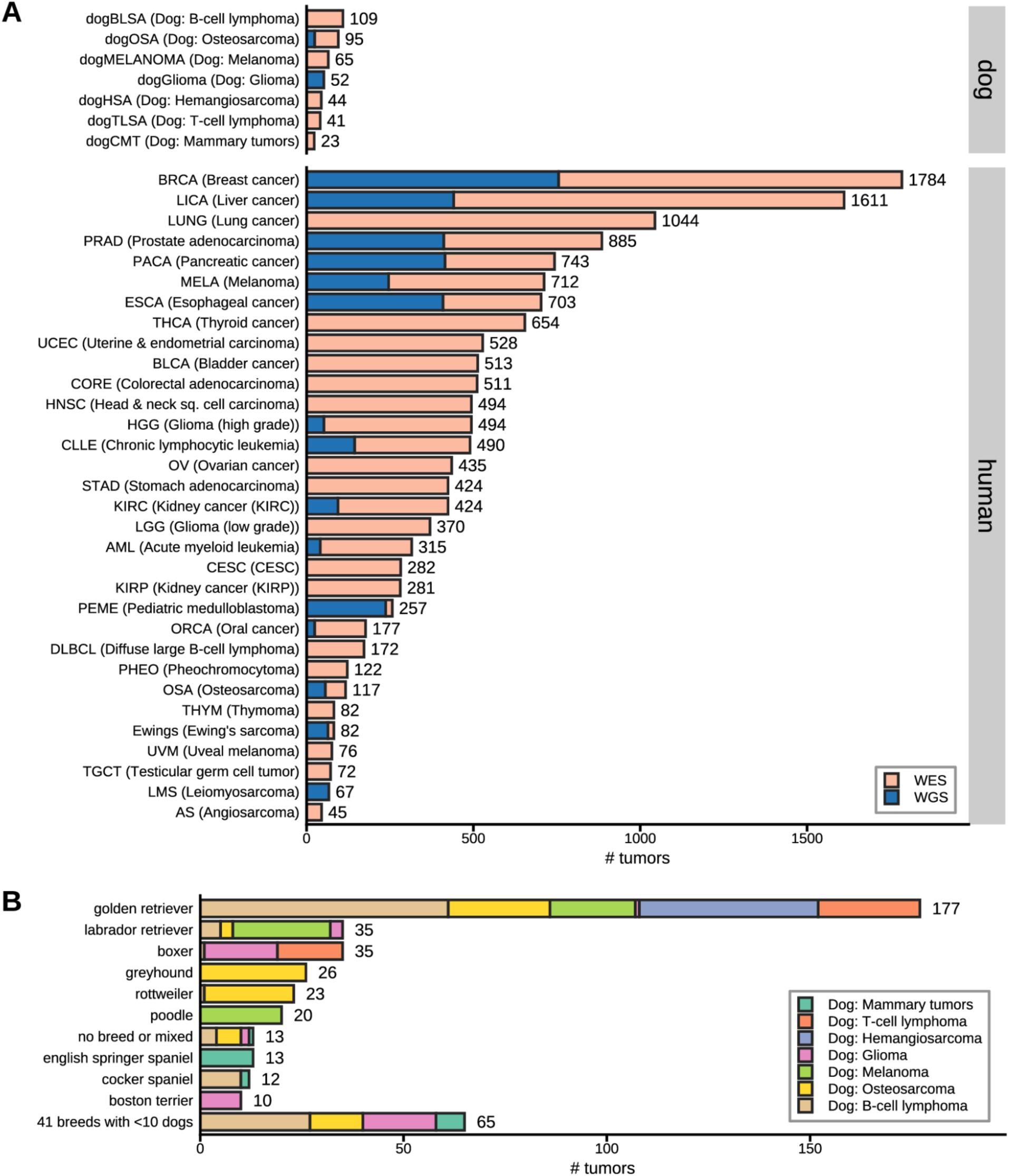
Cancer types and breeds represented in the comparative pan-cancer data set. (**A**) To construct our comparative data set, we compiled existing, publicly available somatic mutation data from whole genome (blue) and whole exome (orange) sequencing from 429 dog cancers and 14,966 human cancers of multiple histologies. **(B)** Dog tumors are from multiple dog breeds, most of which contributed tumors of multiple cancer types.

**Figure S2.**
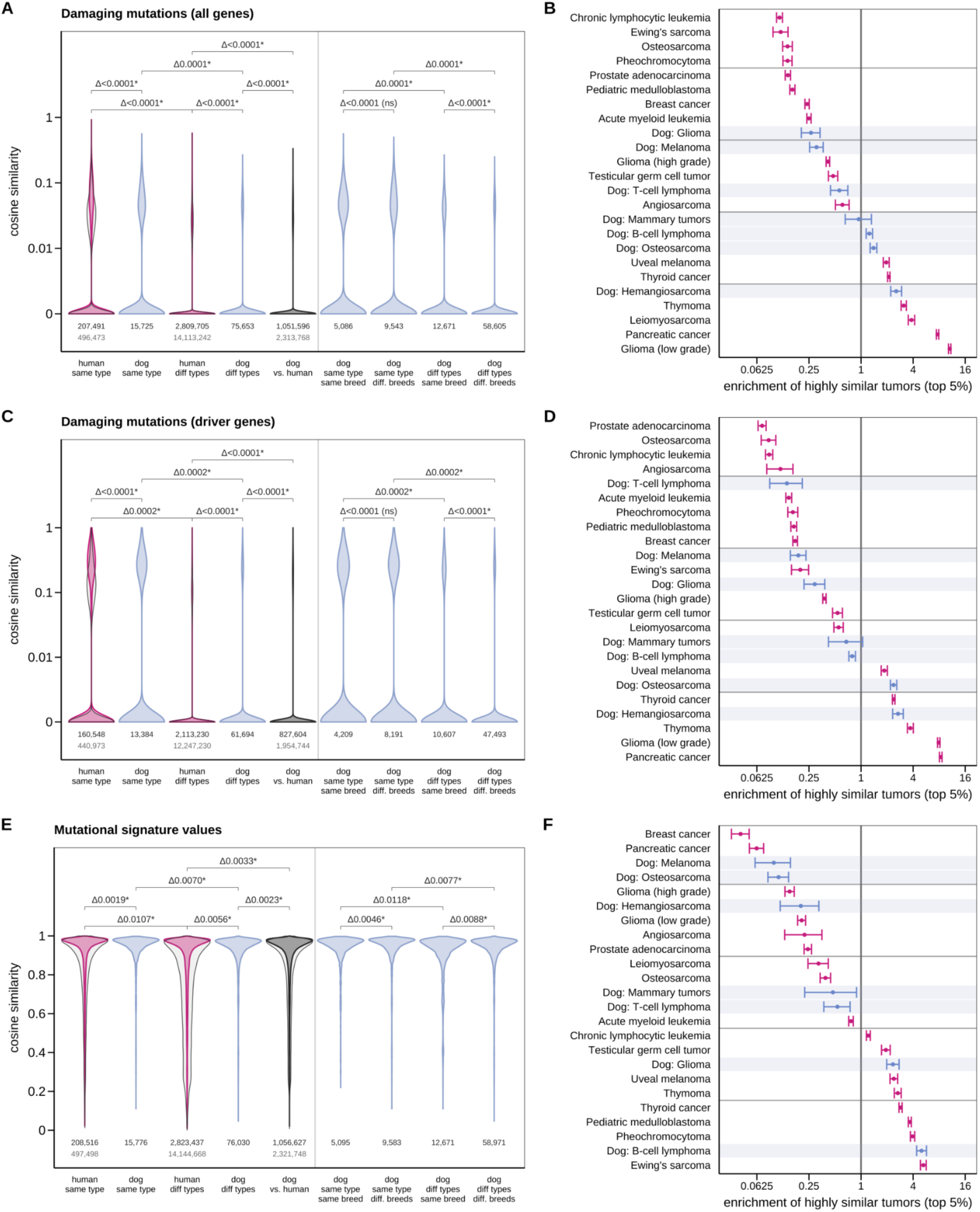
Dog and human tumors are, on average, no more different than dog/dog or dog/human tumors pairs of different cancer types. Cosine similarity scores pairs for pairs of tumors from human (pink) dog (blue), and for dog/human comparisons (black) based on (A) mutations in all genes, with (B) dog cancer types interspersed with human cancer types. The same is seen for (C,D) mutations in driver genes; and (E,F) mutational signatures. Black lines in violin plots show the distributions when the high mutation rate cancer types are not removed.

**Figure S3.**
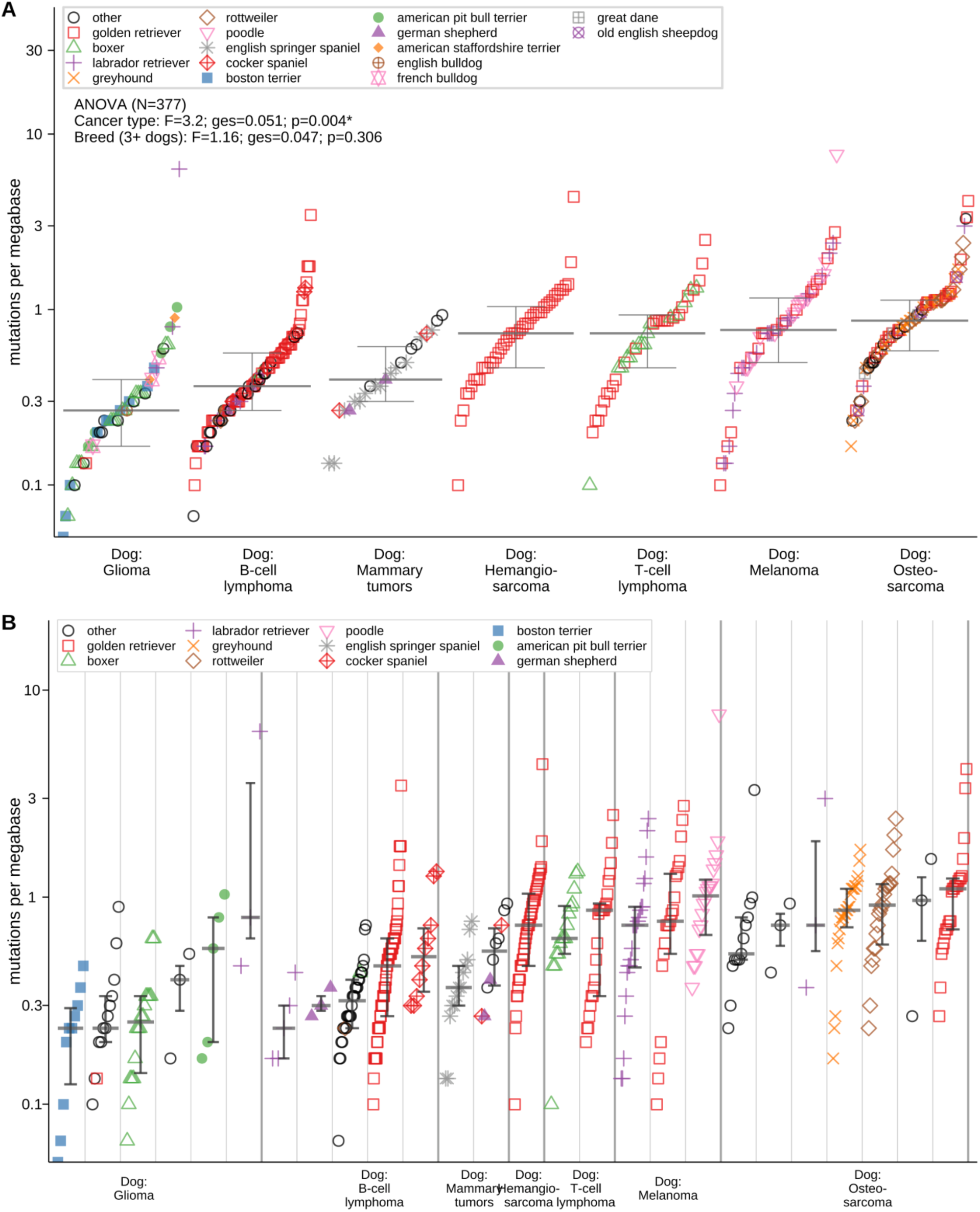
Mutational burden in canine tumors varies between cancer types but not between breeds, Distribution of mutational burden in mutations per megabase **(A)** across all canine cancer types, and **(B)** separated by breed and cancer type.

**Figure S4.**
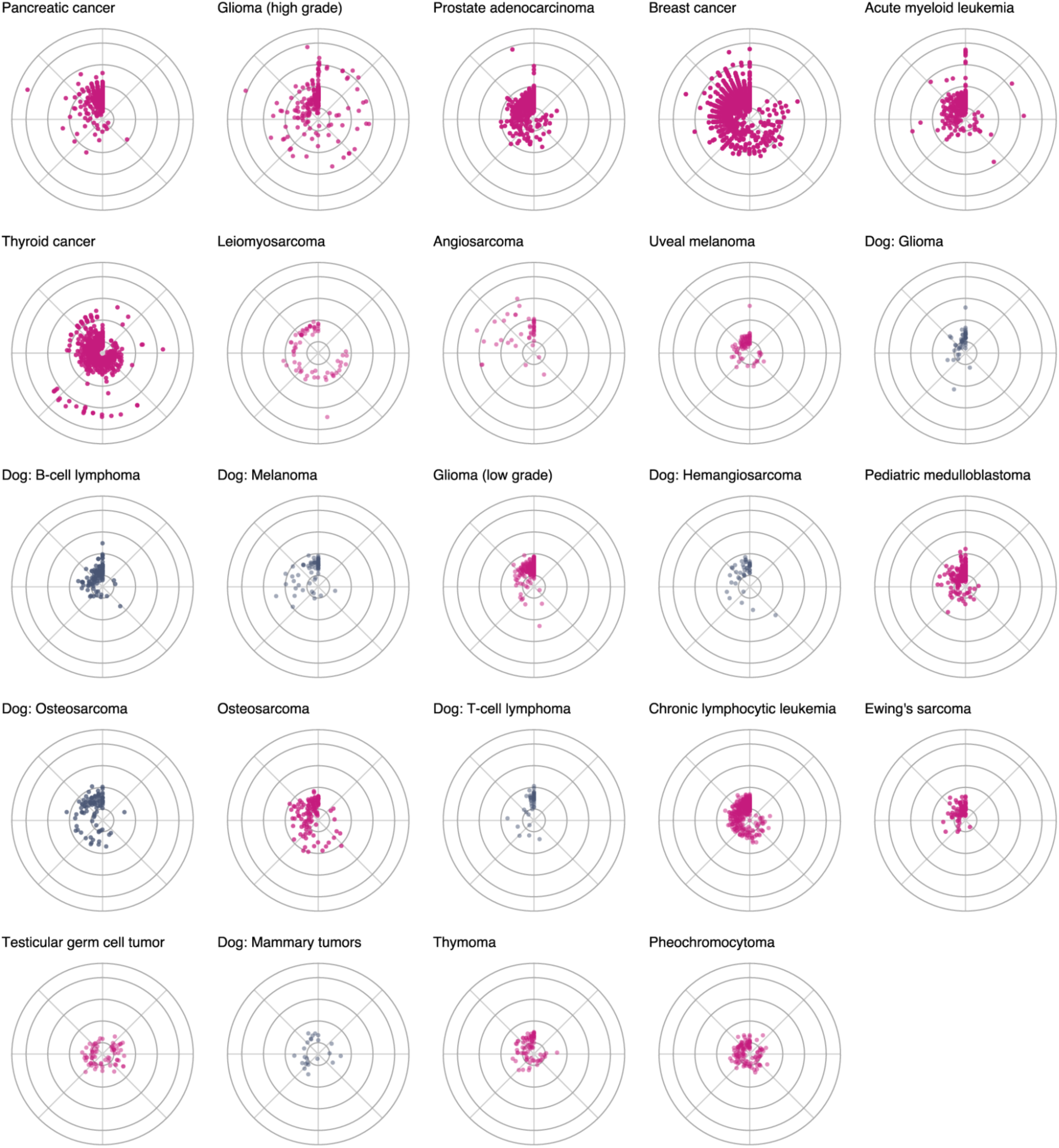
Relative abundance of mutation signatures is heterogeneous within individual cancer types of dog and low-mutation human cancers. Radial plots of each tumor from dog (pink) and human (gray) positioned an angle determined by the relative abundance of 43 mutational signatures detected in at least one human and one dog. Distance from center is log10 of mutation count. Cancer types are sorted by their maximum mutation count.

**Figure S5.**
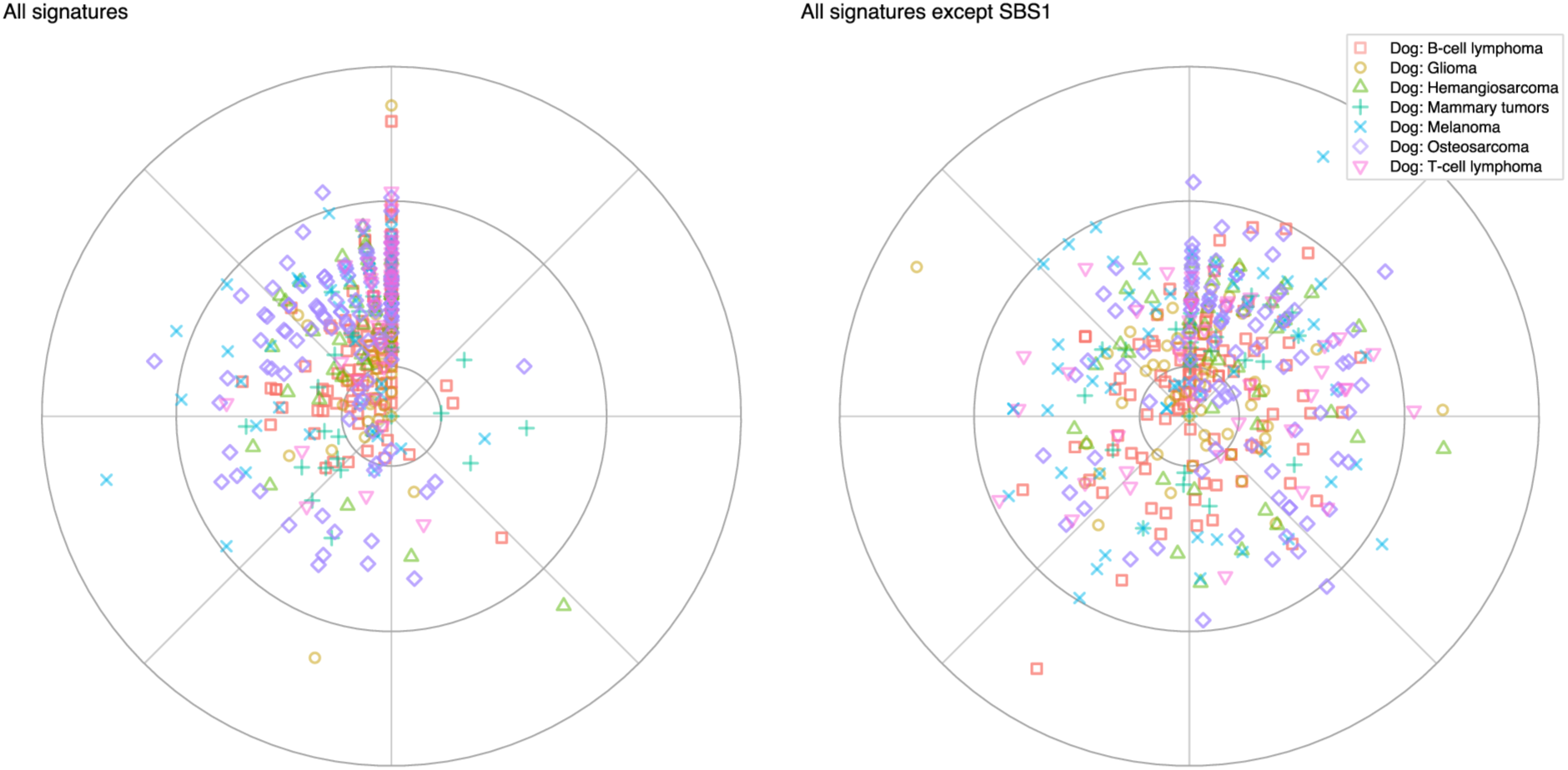
Relative abundance of mutation signatures is heterogeneous within individual dog cancer types. Radial spectrum plots place each tumor into one of 43 (left) or 42 (right, excluding SBS1) sectors, positioned based on its most abundant mutational signature. The angle within each sector reflects the relative contributions of the remaining signatures, and the distance from the center represents the log10 of the mutation count.

**Figure S6.**
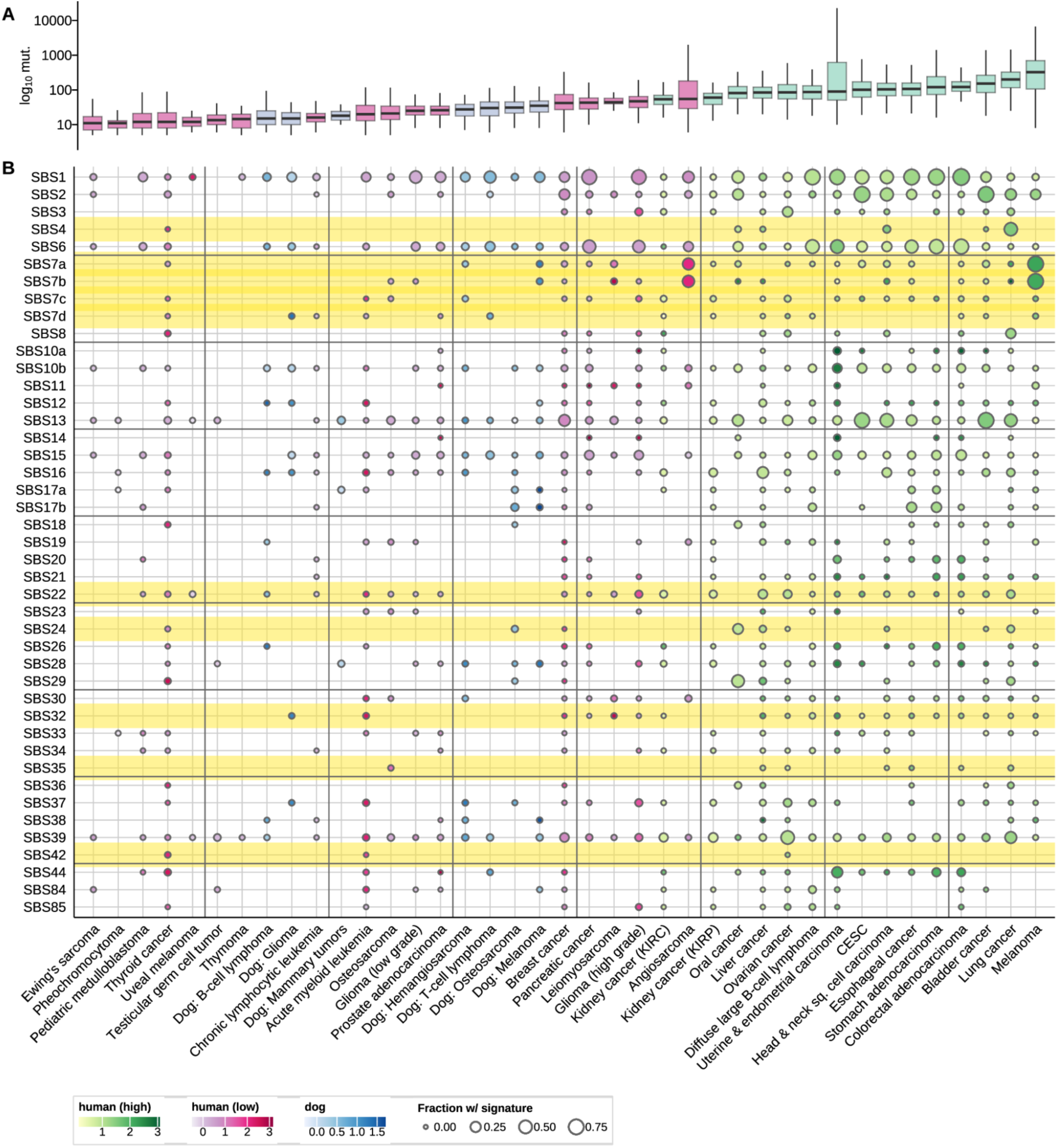
Full version of. figure 4C. Signatures detect in dog and human (both low and high mutation rate) cancer types. Boxplot shows the mutation count per tumor. Circle plot shows how frequent each signature was detected at a value >0.05 in each tumor type. Yellow shading indicates signatures putatitively connected to environmental exposures.

**Figure S7.**
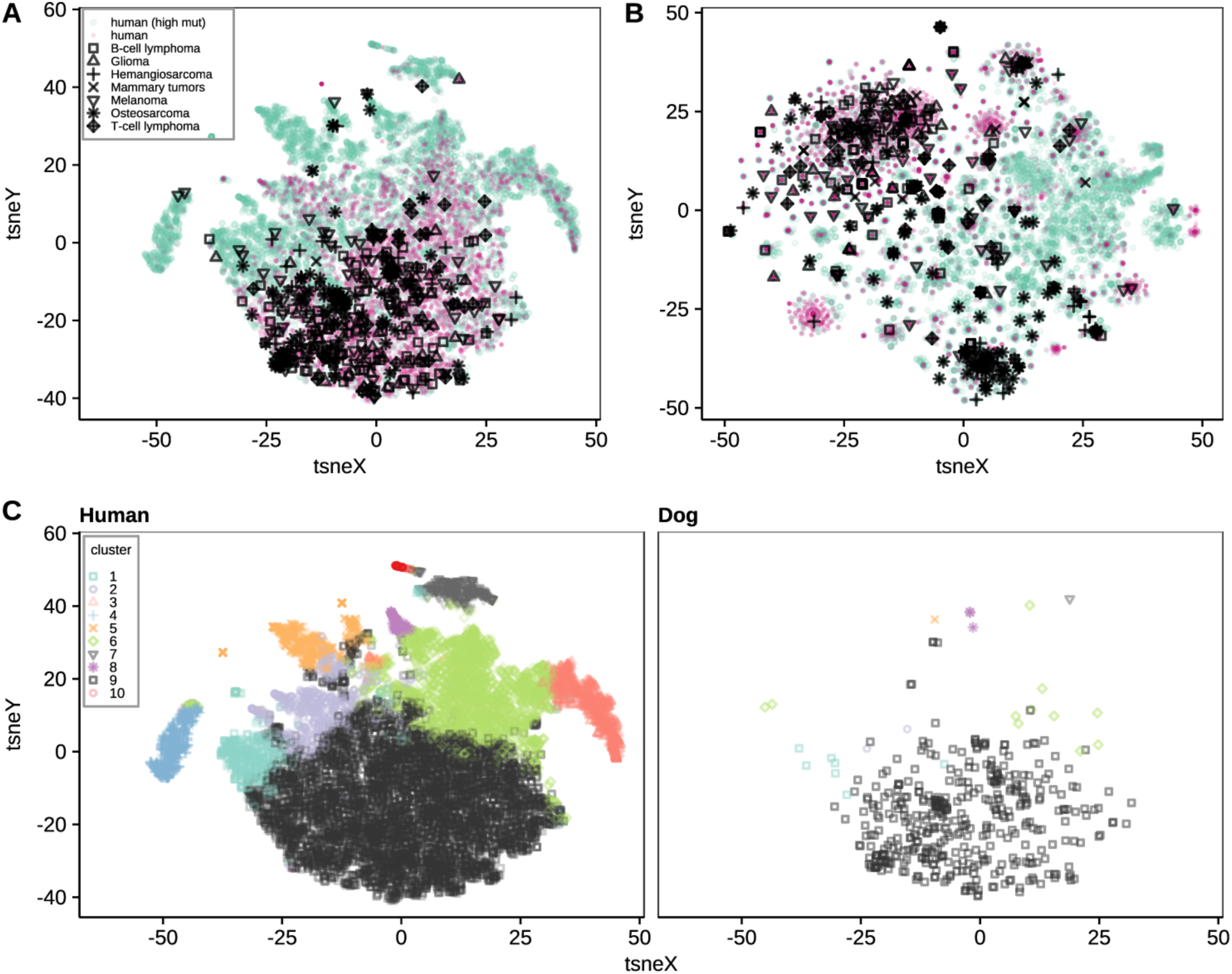
Unsupervised clustering of all dog and human cancer types groups dog tumors with human tumors from low-mutation cancer types. **A.** t-SNE visualization of all dog and human tumors clustered using k-means based on mutational signatures, after dimensionality reduction to principal components explaining 70% of the variance. **B.** Clustering of all dog tumors and human tumors using binary mutation profiles of driver genes. **C.** Joint clustering of all human (left) and dog (right) tumors based on mutational signatures. The position of each tumor is identical to panel A. Cluster 9 contains mostly dog tumors and tumors from low-mutation human cancer types. Similar plots for just the low mutation rate human cancer types and the dog cancer types are in Figure 5.

**Figure S8.**
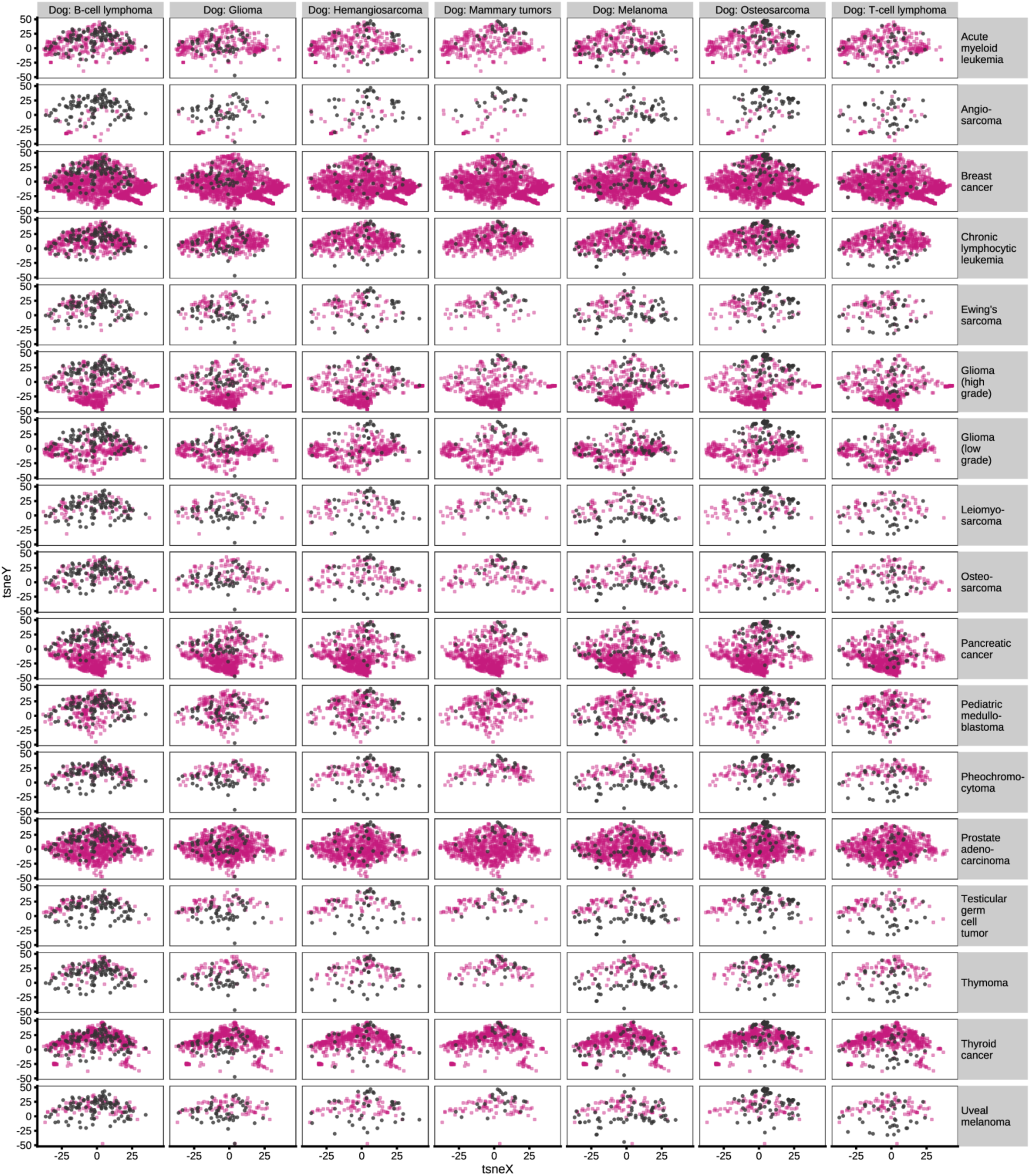
Dog and human cancer types do not form one-to-one matches using mutational signature principal components. Dog tumors cluster with multiple human cancer types and vice versa. Pairwise overlaps are shown for all dog cancer types (black) and low-mutation human cancer types (pink) using k-means clustering on mutational signatures, based on principal components explaining 70% of the variance.

**Figure S9.**
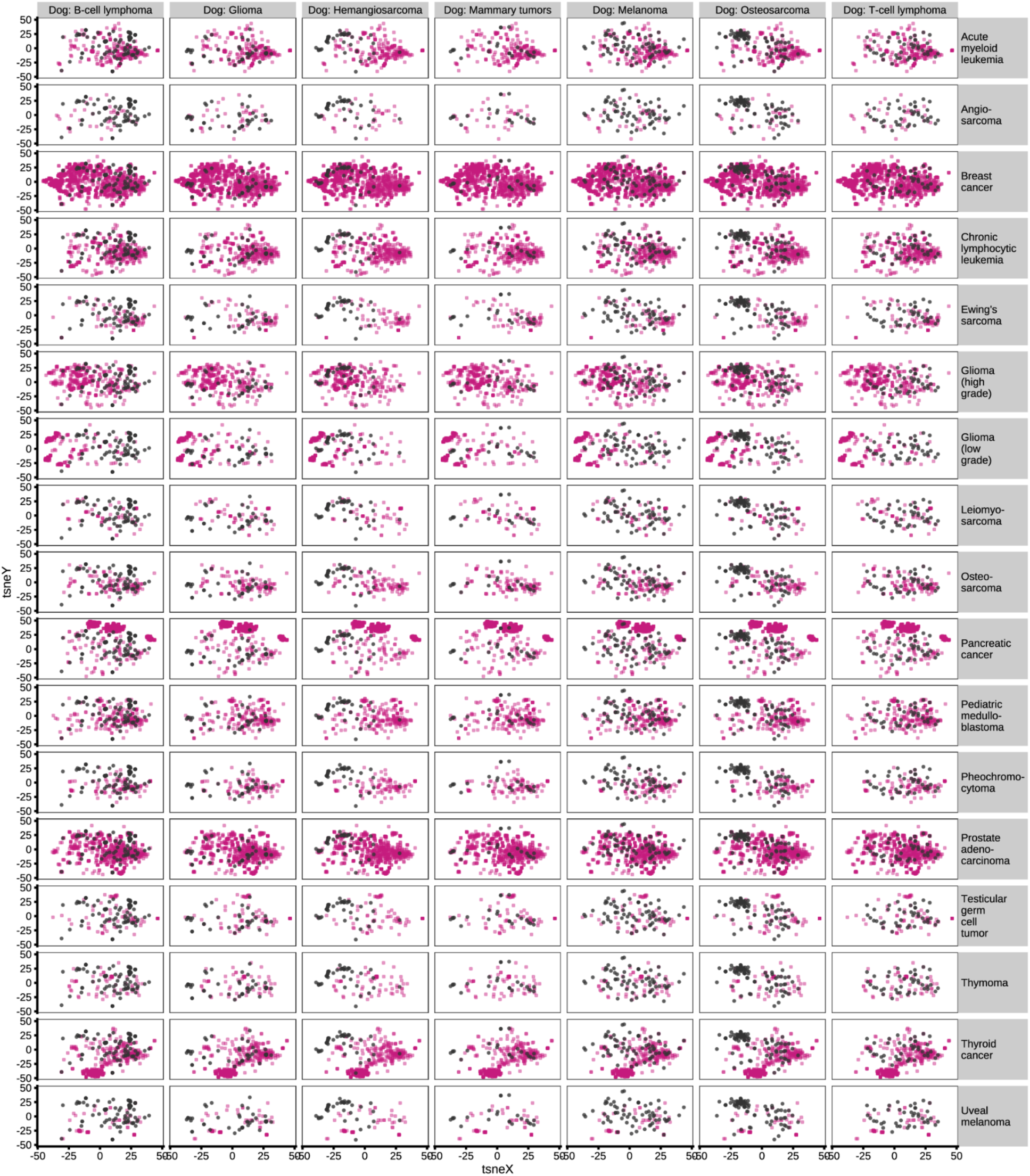
Dog and human cancer types do not form one-to-one matches using driver gene mutations. Dog tumors cluster with multiple human cancer types and vice versa. Pairwise overlaps are shown for all dog cancer types (black) and low-mutation human cancer types (pink) using k-means clustering on driver gene mutations.

**Figure S10.**
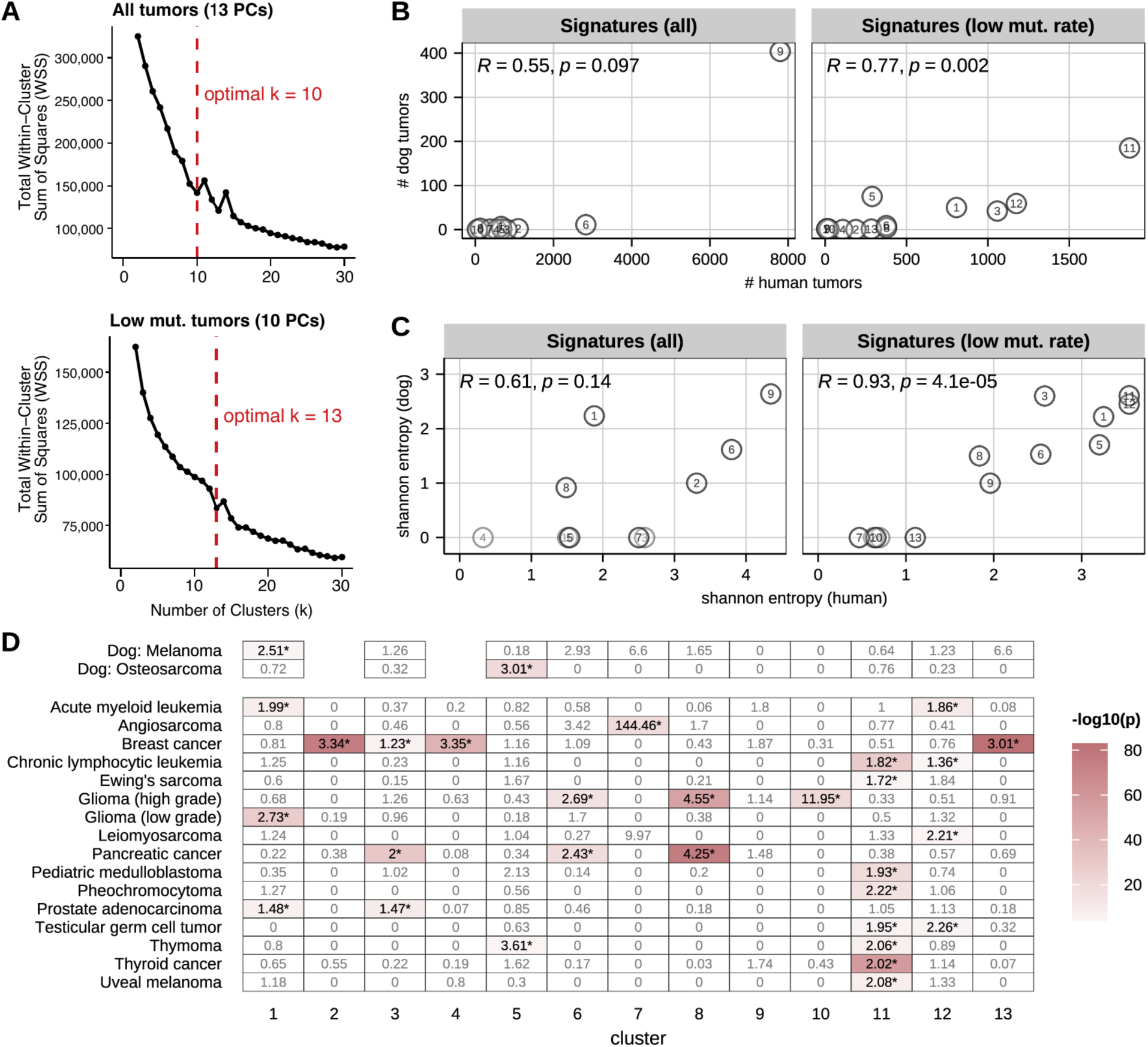
k-Means Unsupervised clustering of dog and human tumors based on principal components explaining 70% of the variance in mutational signatures. **A.** Elbow plot showing the total within-cluster sum of squares (WCSS) as a function of the number of clusters (k) used in k-means clustering. The red dashed line marks the “elbow” point at which adding more clusters yields diminishing returns, suggesting an optimal number of clusters. **B.** When clustering all tumors, most tumors fall in cluster 9. Removing high mutation rate human cancer types yields more clusters with more tumors in both species and **C.** more heterogeneous clusters, as measured with Shannon’s entropy. **D.** Tumors of some cancer types are enriched in some clusters compared to the null expectation of random distribution.

**Figure S11.**
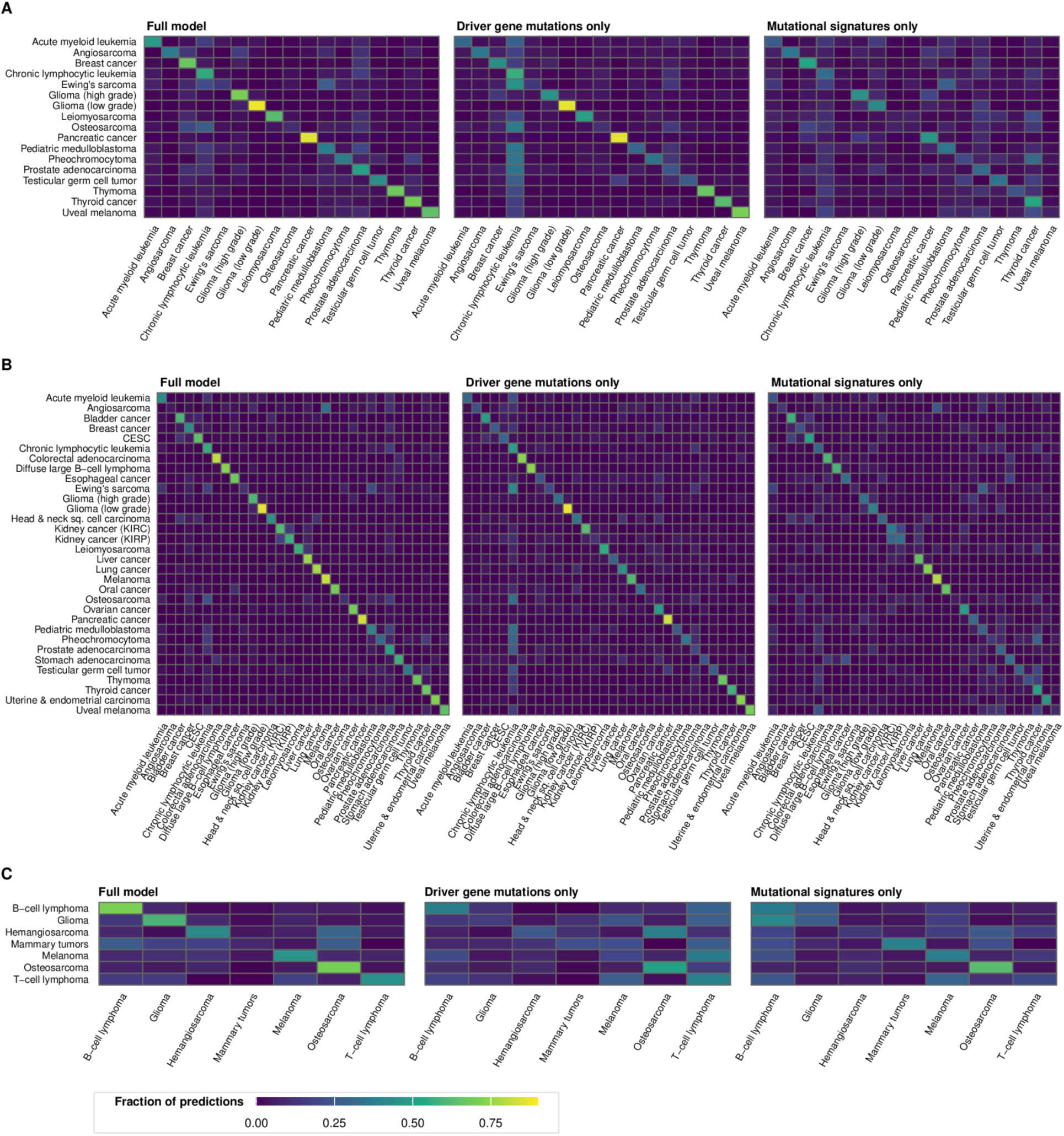
Confusion matrices. Confusion matrices depicting the proportion of tumors of each cancer type (y-axis) classified as each cancer type (x-axis) for each machine learning model. **(A)** Human trained classifiers on low mutation rate human cancer types only. **(B)** Human-trained classifiers on all human cancer types. **(C)** Dog-trained models on all dog cancer types.

**Figure S12.**
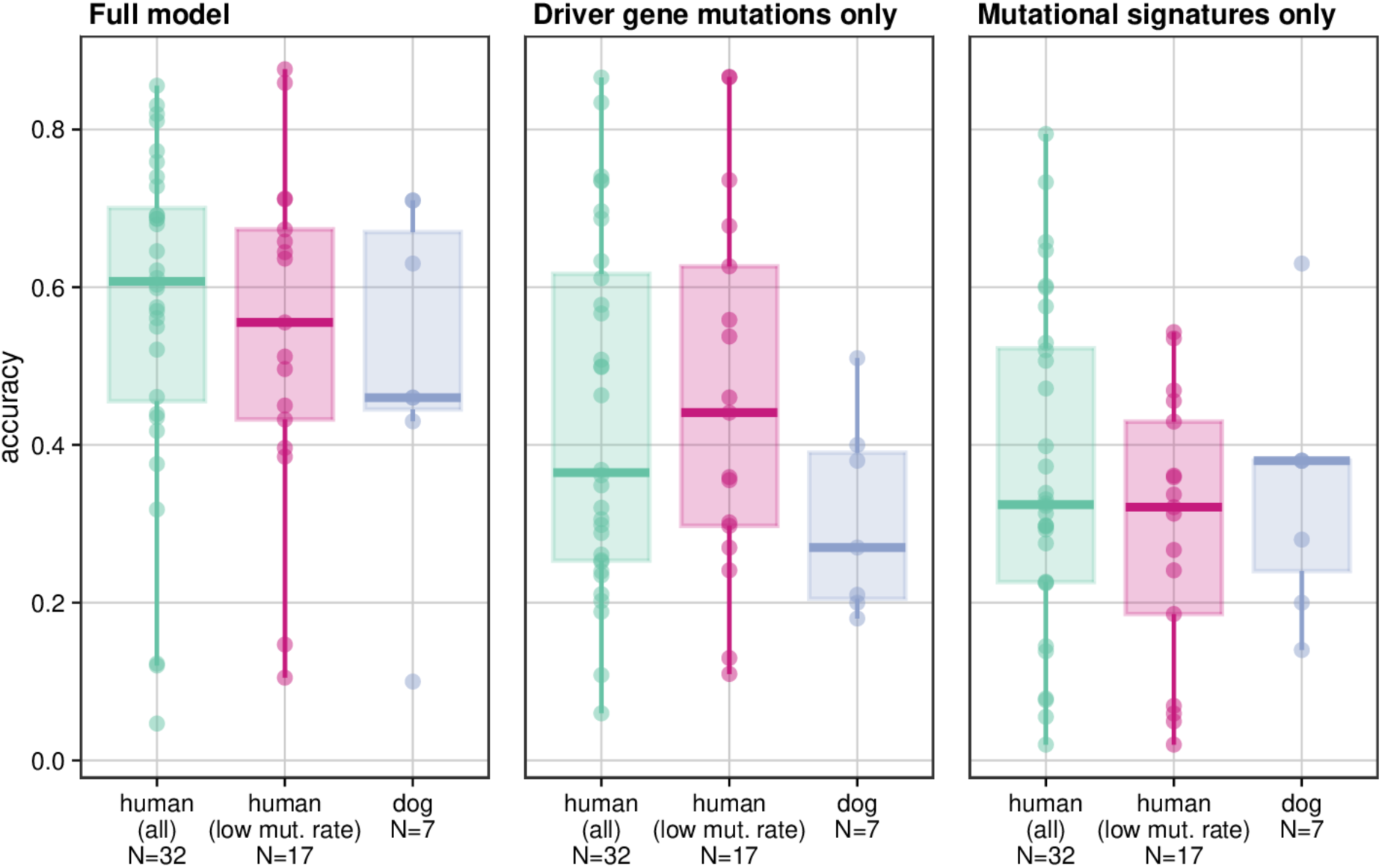
Average accuracy per cancer type. Boxplots depict the average accuracy across five folds for each cancer type in each machine learning model. Green - human-trained, all cancer types; Pink - human-trained, low mutation rate cancer types; Blue - dog-trained.

**Figure S13.**
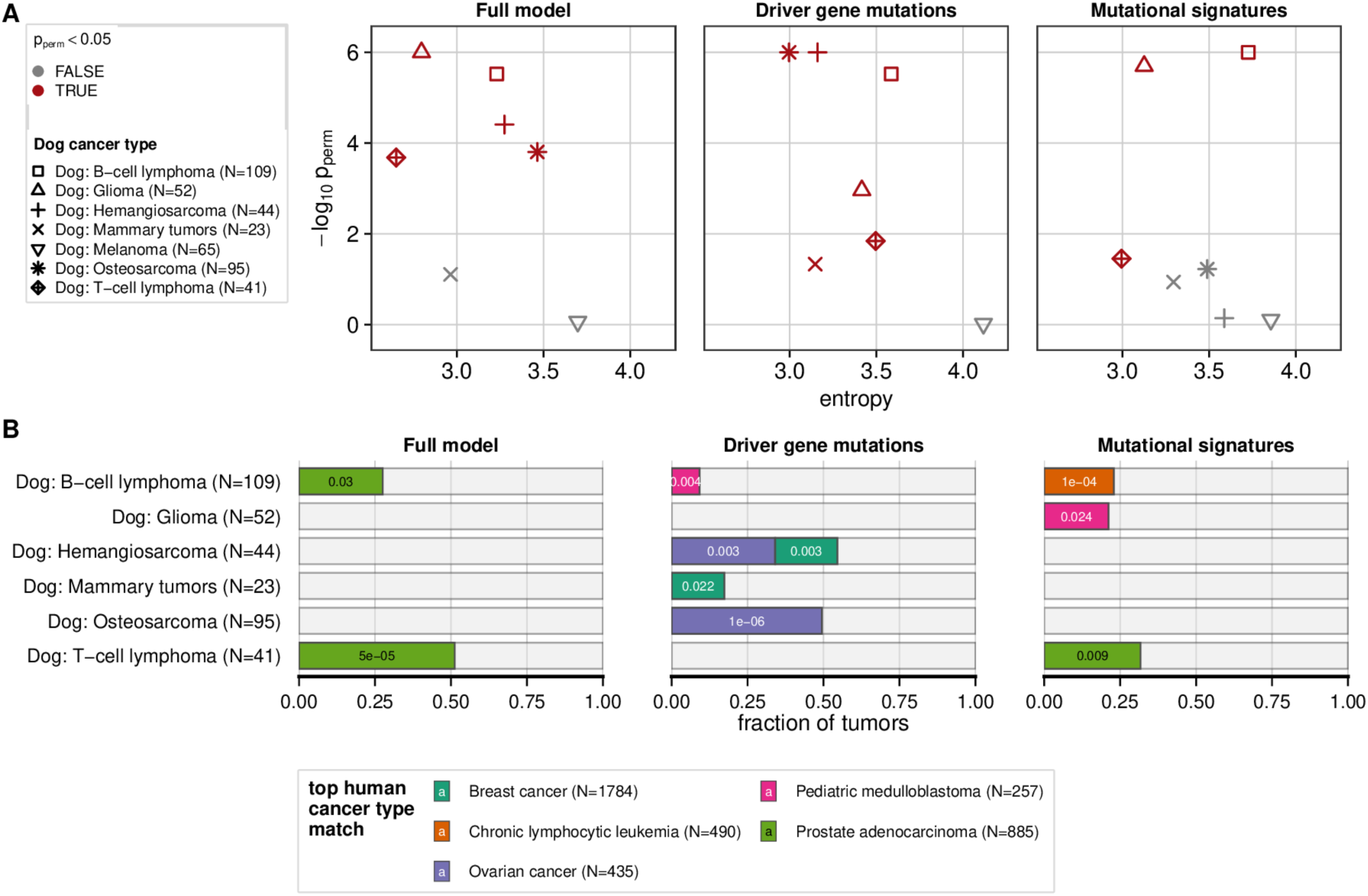
Clumpiness and one-to-one cancer type matching in the human-trained, all cancer types models. (A) Dog cancers in red were classified as fewer human cancer types than expected by chance. (B) A subset of dog tumors were classified as specific human cancer types more than expected by chance in each model.

**Figure S14.**
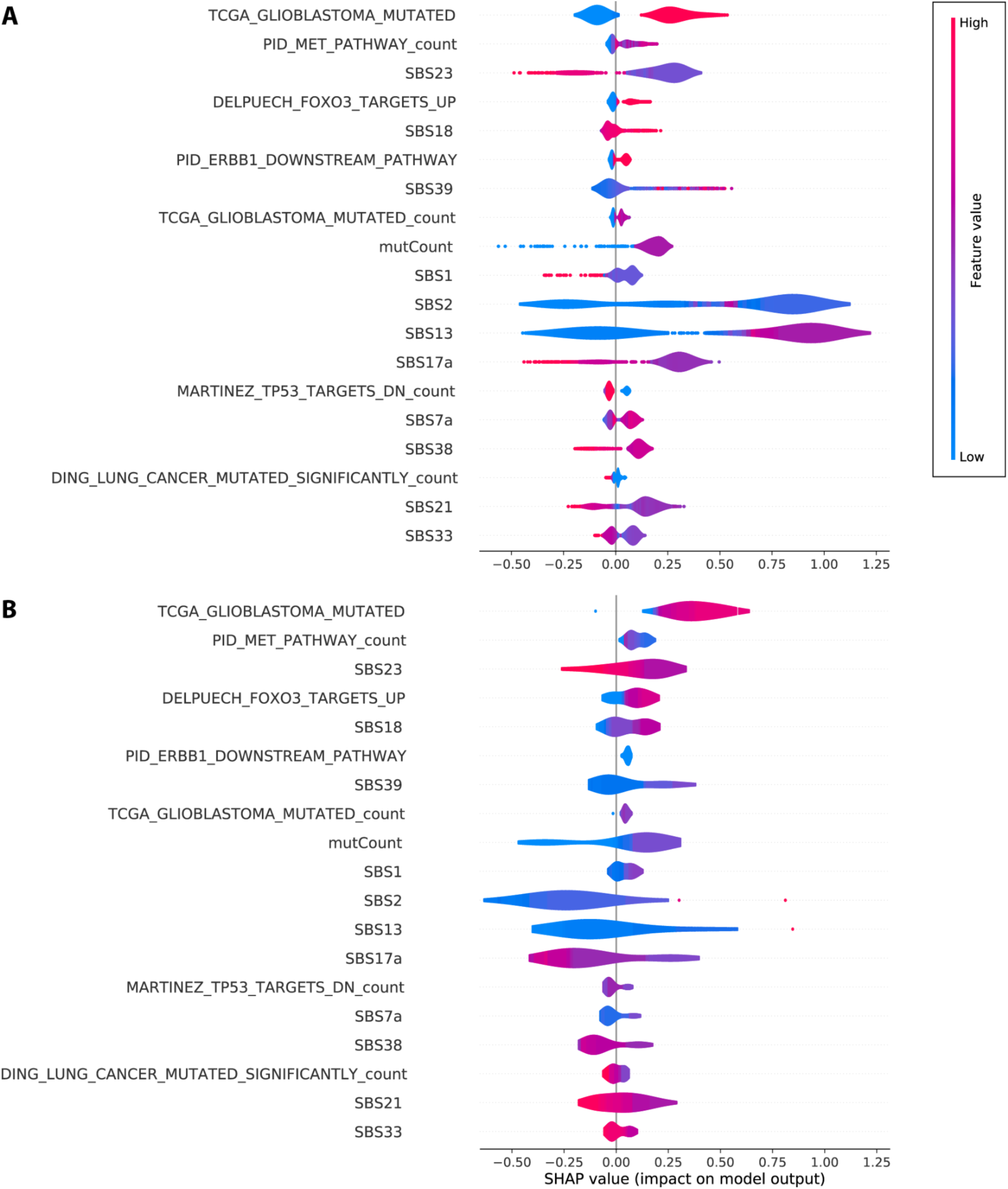
Shapley and feature value violin plots (breast cancer). The top and bottom ten most important features for classification of dog hemangiosarcoma tumors as breast cancer, visualized in (A) human breast cancer tumors classified as breast cancer, and (B) dog hemangiosarcoma tumors classified as breast cancer.

**Figure S15.**
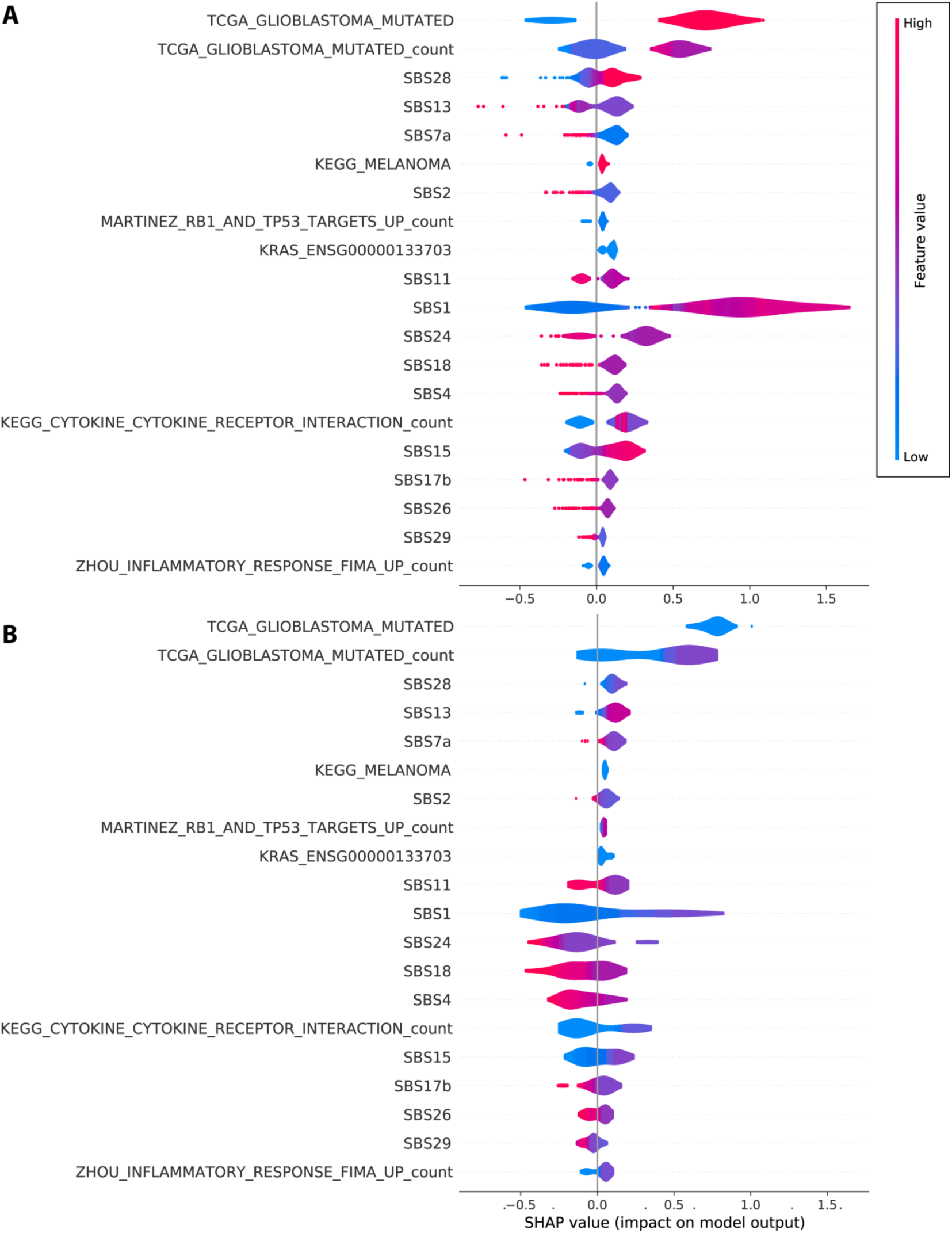
(HGG shapley) Shapley and feature value violin plots (high-grade glioma). The top and bottom ten most important features for classification of dog hemangiosarcoma tumors as high-grade glioma, visualized in (A) human high-grade glioma tumors classified as high-grade glioma, and (B) dog hemangiosarcoma tumors classified as high-grade glioma.

**Fig S16.**
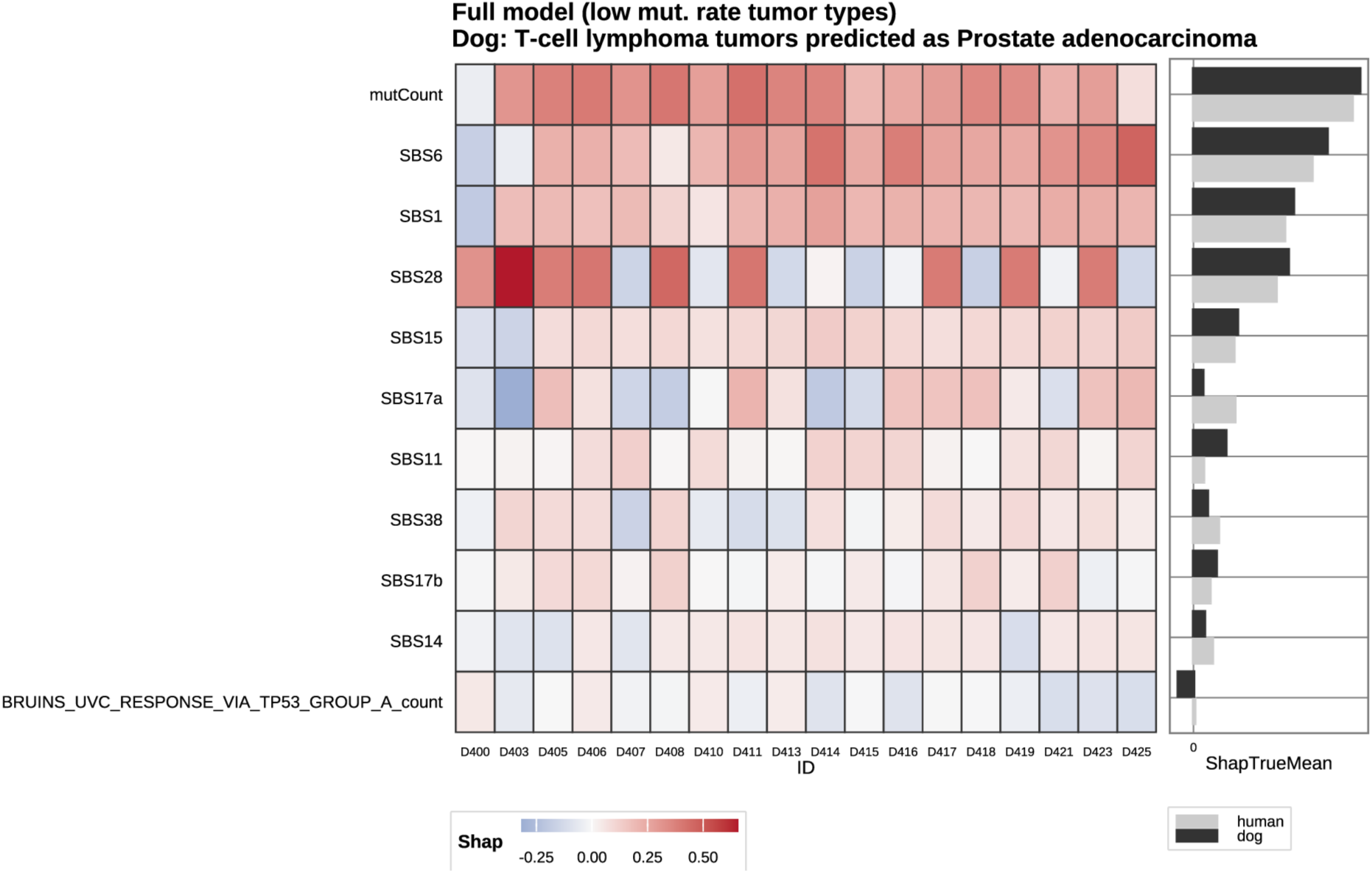
TLSA Heatmap of Shapley values (dog T-cell lymphoma classified as prostate adenocarcinoma). The top and bottom most important features for classification of dog T-cell lymphoma tumors as prostate adenocarcinoma, by tumor.

**Fig S17.**
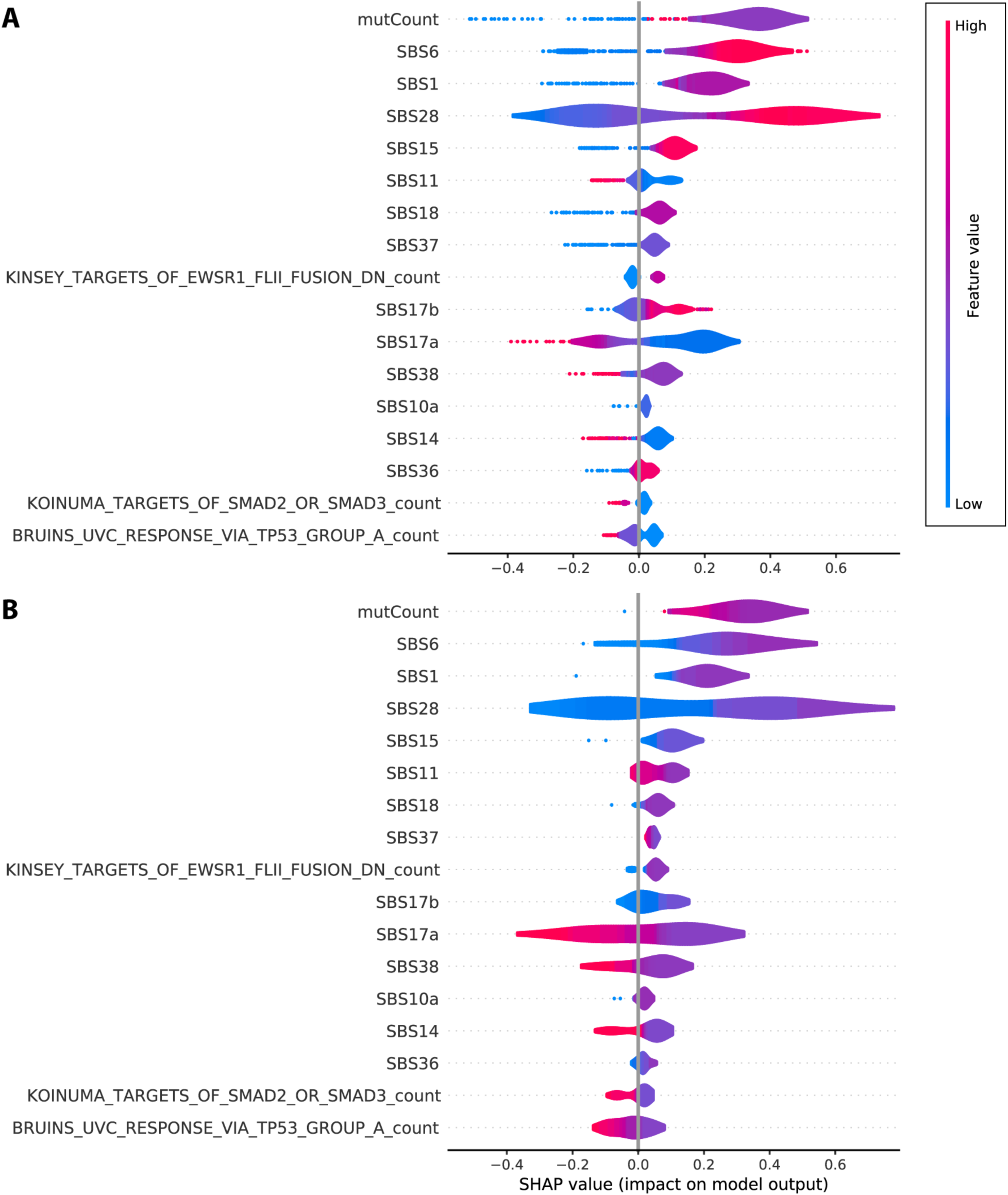
Shapley and feature value violin plots (prostate adenocarcinoma). The top and bottom ten most important features for classification of dog T-cell lymphoma tumors as prostate adenocarcinoma, visualized in (A) human prostate adenocarcinoma tumors classified as prostate adenocarcinoma, and (B) dog T-cell lymphoma tumors classified as prostate adenocarcinoma.

## Supplementary Methods

### Acquisition of human data

We obtained publicly available human somatic mutation data from The Cancer Genome Atlas (TCGA), the International Cancer Genome Consortium (ICGC)^1^, and cBioPortal^2^. We downloaded data from the 33 projects in the TCGA program on the Genomic Data Commons (GDC). Individual study data from the ICGC Data Coordination Center (DCC) Data Release 28 (excluding TCGA studies) were downloaded from the DCC Data Releases page. In addition, we downloaded data from individual studies from cBioPortal. Data sources are listed in **<u>Table S1</u>**.

### Acquisition of dog data

We compiled somatic mutation data from WES and WGS sequencing studies of seven canine cancer types and subtypes. We leveraged mutational calls from previously published studies or preprints. For B-cell lymphoma^3,4^, T-cell lymphoma^3^, osteosarcoma^5^, and hemangiosarcoma^6^, we downloaded published mutational data for osteosarcoma^7^, and glioma^8^. Mutational calls were provided pre-publication by the authors for oral melanoma^9^ and mammary tumors^10^. NCBI, SRA, or ENA sample identifiers are provided in **Table S19**.

### Cancer data sets included

We included all studies from the TCGA and ICGC data sets that considered a single cancer type. For studies that contained data from multiple cancer types, we separated samples by cancer type when feasible, otherwise, the entire study was excluded. To include samples for the human cancer types for which we had canine data from the histology-matched human cancer (angiosarcoma, osteosarcoma, and diffuse large B-cell lymphoma), or to increase the number of samples included, we also included data from studies hosted on cBioPortal.

### Data QC

We applied consistent criteria in compiling published data from multiple sources. Overall, we favored data from primary tumors rather than metastatic sites, and from earlier-rather than later-stage cancers. We also sought to include no more than one sample from any individual patient. The metadata types available differed somewhat among sample sources, so we filtered as follows:

#### Human cancer QC

##### cBioPortal studies

We removed samples marked as being from relapse time points, as well as those that, based on ID, were duplicates of samples in the included ICGC studies. Single angiosarcoma samples per patient were selected as a subset of the data_clinical_sample.txt file provided with the study data. The first available time point (based on the BX_TIME_DAYS column, annotated as “Sample collection time in days from primary diagnosis”) was selected for each patient, regardless of site or procedure. In one case, two samples labeled T1A and T1B were both annotated from the same time point, so the first listed (T1A) was selected. Samples from the Columbia University pediatric study (study identifier: mixed_pipseq_2017) were selected if “osteosarcoma” appeared in the “Cancer Type Detailed” column, “WES” appeared in the “NGS Test” column, and “Diagnostic” appeared in the “Sample_Timepoint” column. Ten samples were found to be shared between the DFCI (study identifier: dlbcl_dfci_2018) and Broad Institute (study identifier: dlbc_broad_2012) DLBCL studies. The data for these samples from the DFCI study was retained, and the corresponding data from the Broad Institute was excluded.

##### ICGC studies

We excluded samples that did not have either whole genome or whole exome sequencing data and/or were not from a primary or metastatic tumor sample. When more than one sample was present from a given patient, we prioritized (in order of importance): solid tissue primary tumors to other types of primary tumor; primary tumors to metastatic tumors; if listed, higher percentage cellularity to lower; whole genome sequencing to whole exome; if listed, earlier BINET grade to later BINET grade; enrichment by flow sorting to enrichment by laser capture microdissection; and samples included in the Pan-Cancer Analysis of Whole Genomes over not. For most ICGC sample sets, we excluded all samples if multiple samples remained following application of these criteria. Two studies, LMS-FR and UTCA-FR, contained samples from cancer types for which few samples were available from other studies, so we retained one randomly selected sample per patient.

#### Dog cancer QC

We excluded eight samples from the canine melanoma dataset because they were outliers for the number of somatic mutations called. The average mutational count for the remaining 69 samples was 465 (range: 136-1435). One excluded sample had a single somatic mutation. The other seven excluded samples had mean 36125 mutations (range: 11652-56641). Glioma samples were excluded if whole genome or whole exome sequencing was not available or if there was no matched normal. An additional sample was excluded due to being annotated as not having tumor content. Twenty-three of the canine mammary tumors were selected for our analysis based on histology (including carcinomas and mixed tumors, while excluding hyperplasia, adenoma, fibrolipoma, other cancer types (*e.g.* chondrosarcoma), and samples without histology data.

### Data harmonization and processing

We defined a set of features for comparison across cancer types and species.

#### Identifying genes orthologous between human and dog

We used Ensembl Biomart and NCBI Gene^11^ databases to identify a set of 15,315 orthologous genes to compare between human and dog. In cases where the two sources disagreed in defining a given ortholog, all genes identified by either source were considered orthologs of one another. To define the start and end positions of the coding region for each gene, we used annotations from Ensembl (version 104 for canFam3 and hg38; version 87 for hg19). Genes that were predicted to have a length of zero in either species were removed.

#### Defining known cancer-driver genes

From among the 15,315 orthologous genes, we defined a subset of 1,098 known or suspected driver genes from the Catalog of Somatic Mutations in Cancer (COSMIC) Gene Census^12^, as well as genes known to be affected by clonal hematopoiesis in humans or commonly affected in canine cancers. We also designated 4,852 genes that are targeted by existing drugs based on the OncoKB database^13^ (**<u>Table S3</u>**).

#### Identifying predicted-damaging variants

Variants were parsed from Variant Call Format, Mutation Annotation Format (MAF), or custom formats using a custom python script leveraging the pysam^14^ wrapper of the Samtools^15,16^ and htslib^17^ tools. We included in our analysis variants that fell within the coding regions of our 15,315 orthologous genes and that were predicted to be damaging. In the human datasets, “masked” variants where both reference and alternate alleles were set to reference were removed. We defined as damaging variants that were classified as moderate or high impact by SnpEff, or those that had ontologies in Variant Effect Predictor (**<u>Table S18</u>**).

#### Defining mutated genes and pathways

To define mutated genes, we opted for a binary classification of damaged or not damaged. We used the output of VEP or SNPEff, as available, to determine whether or not a gene was damaged in a sample. We used this same concept to define mutated pathways. Using the C2 database of curated pathways from MSigDB, we defined a pathway as mutated if any of its constituent genes was damaged. We removed from the analysis any samples that had fewer than five mutations remaining after data harmonization.

#### Defining and assessing hot spot mutations

We defined mutational hotspots at the codon level, considering a mutation in any of the three bases of a given codon to be a hit to that codon. To construct a set of known hotspots for which to tally mutations, we used sites listed in the Database of Curated Mutations (DoCM)^18^ and from Hess, *et al*.^19^, using a cutoff of Q_LNP < 0.1. We also considered as a hotspot any site mutated in five or more of our samples. To enable consideration of the same codons across all samples, we used UCSC’s liftOver tool to liftover variants from samples aligned to canFam3 and hg19 samples to hg38.

#### Calling mutational signatures

We used SigFit v.2.2^20^ to quantify mutational signature exposures for each sample We tallied trinucleotide context mutational opportunities in the orthologous regions included in our data across hg19, hg38, and CanFam3.1 using the Biostrings package v.2.62^21^. We used mutations observed in each sample, along with the relevant reference genome, to generate trinucleotide context mutation counts for each sample. We fit all of the samples’ trinucleotide counts to the normalized COSMIC v3 single base substitution (SBS) signature set^22^ using 100,000 iterations and 50,000 warm-up iterations, normalizing each sample by its genome’s mutational opportunities. We retained all signatures that were significantly different from zero (lower bound estimate was at least 0.01, the default value for SigFit v.2.2) in at least one sample and refit the samples using only those signatures. To calculate signature values for each species as a whole, we performed the same process as above using the sum of the trinucleotide counts across all samples in that species as the input.

##### Determining granularity of cancer types

Several of the human cancer types represented in our set (*e.g.,* breast cancer, uterine, and endometrial cancers) have multiple known subtypes. To determine at what level of granularity to group the samples, we assessed classification accuracy under machine learning models trained on human data, using extreme gradient boosting (XGBoost). If classification accuracy for a given subtype was < 0.25, it was removed. If a subtype within a given cancer type had classification accuracy ≥ 0.25 but lower than for other subtypes within that same cancer type, it was either made its own category or removed. Specific samples that could not be classified into the resulting categories were excluded from further analysis.

##### Defining low-mutation cancer types

To define a set of human cancer types with mutation rates comparable to those observed in dogs (“low mutation rate human cancer types”), we applied a conservative threshold of excluding any human cancer type whose median mutation rate exceeded that of the most highly mutated dog cancer type (dog osteosarcoma; median = 0.43 mutations/Mb). However, to enable comparisons between histologically matched and unmatched cancer types, we included two human cancer types which are histologic counterparts to dog cancers and have slightly higher mutation rates: high-grade glioma (0.57 mutations/Mb) and angiosarcoma (0.62 mutations/Mb). We did not include the two histological matches that were amongst the most mutated human cancer types: diffuse large B-cell lymphoma (1.06 mutations/Mb) and melanoma (3.14 mutations/Mb).

##### Computing pairwise similarities of samples

To quantify pairwise genomic similarity of tumors, we used the cosine function as implemented in the R package *coop*. Cosine similarity scores can range from 0 (indicating that the tumors are maximally distinct) to 1 (indicating that the two tumors are identical). We computed cosine similarity scores using both damaging mutations and signature values as input.

##### Computing angles for radial spectrum plots of signature values for each tumor

To display signature values and mutation counts for each tumor, we used radial spectrum plots, using an approach adapted from the methods that Lawrence et al.^23^ introduced for plotting tumors according to the relative abundance of classes of mutations. For each tumor, we computed αngle = 2πΣ ^K^ _r=1_ i_r_(1/K) ^r^. Here, K refers to one of the 43 circle sectors to which a given tumor is assigned, Each sector represents one of the 43 signatures we quantified for each sample. In turn, r indicates the rank for that signature in a given sample. This approach places each tumor within the sector that represents its most abundant signature, and within a series of nested subsectors corresponding to the remaining signatures, by declining rank. Each tumor is placed at distance from the circle equal to log_10_ of its mutations count.

#### Unsupervised clustering

We implemented unsupervised k-means clustering of all dog and human tumors using the kmeans function in stats version 4.4.3 library in R0. To cluster tumors based on signature values, we first inferred principal components, then used as input for k-means clustering the minimal set of components required to explain 70% in sample-wide variance. To cluster tumors based on driver-gene mutation states, we applied k-means directly to binary mutation data, as, for these sparse data, principal components analysis did not reduce the number of features required to capture 70% of total variance. For each data type, we used the elbow method to determine the number of clusters that minimized within-cluster sums of squares. To enable visualization of these high-dimensionality data, we use t-distributed Stochastic Neighbor Embedding as implemented using the Rtsne library in R.

#### Machine-learning classification

We trained extreme gradient boosting (XGBoost) machine-learning models to classify tumors from humans and dogs. We trained models using human data and using dog data for both all cancer types for exclusively low-mutation cancer types. Feature selection. We used a random forest approach for recursive feature elimination with cross validation to select the most informative from among the full set of mutated genes and pathways, mutational signature values, and overall mutation count. This process uses a Bayesian optimization process to repeatedly train models and narrows down optimum values. In each round, this process eliminates 100 features, until a minimum of 400 features are chosen. Model training. We randomly subsetted our data into five folds. For each of these folds, we used 80% of the samples for training the XGBoost model, holding out 20% for testing.

#### Calculating fraction of samples correctly classified

We computed the overall accuracy of each model as the fraction of samples assigned to the correct cancer type in the classifier trained on data from that species.

#### Shapley values

We computed per-sample/per-classification Shapley values to identify the impact of each feature on the probability mass assigned to a given cancer type for a given sample using the (TreeExplainer) function from the (fasttreeshap) package^24^ in python. As input, we used the pruned set of feature types used in making predictions, and the output probabilities per sample per classification.

#### Assessing classification clumpiness for dog cancers

To measure clumpiness—the extent to which the best matches for each dog tumor were concentrated in particular human tumor types—we used Shannon’s entropy, and assessed the significance of clumping by label permutation. We used a permutation-based approach to assess, for samples of each histological label, whether the human- and dog-trained classifier assigned distribution probability mass to a smaller number of cancer predictions than anticipated by chance. To quantify distribution of probability mass for the true samples of a given histological label, we summed probability-mass distributions across samples, then computed the Shannon entropy ( (−1*sum(p_i*ln(p_i)). Larger entropy values indicate greater evenness of density across categories, with maximum entropy equal to the total number of categories.

Therefore, the lower the entropy for a given histological label, the more “clumped” the probability mass assigned by the classifier. To assess the significance of clumping for each histological label, we drew one million sets of samples at random and with replacement from the full sample set, with sample size matched to the true sample size for each histological label. We computed the entropy for each of these random draws. To compute a one-sided p-val for the probability of observing entropy as low as observed for the true samples by chance alone, we computed the fraction of randomized samples that yielded an entropy at least as low as that for the true sample set.

#### Assessing “significantly associated” histological label / classifier label pairs

To assess which histological labels are associated with a particular classifier label more than expected by chance, we again used a permutation test. For each histological label, we first summed the probability weights outputted by the classifier grouped by classification label to get a cumulative probability sum for each histological label / classifier label pair. For each histological label we then generated one million sets of samples at random and with replacement from the full sample set for that species, where the size of each sample was equal to the number of samples with that histological label. For each of these sets, we computed the cumulative probability sums as we did for the original set and retained the maximum sum for that set. To compute a one-sided p-value for the probability of a given histological label being associated with a given classification label, we calculated the fraction of sets drawn for that histological label whose maximum sum is at least as high as the sum for the histological label / classification label pair.

